# STR-PG: A Topology-decoupled Pangenome Framework for Scalable Short-read Genotyping of Short Tandem Repeats

**DOI:** 10.64898/2026.08.07.743532

**Authors:** Jiajing Yuan, Zhengfa Xue, Haoran Tang, Yuqian Liu, Jiayin Wang

## Abstract

Short tandem repeats (STRs) are a rich and highly polymorphic source of human genetic variation, but representing and genotyping them in pangenome graphs remains challenging. Explicitly encoding each STR allele as a separate graph path results in increasingly complex local structures as cohort diversity increases, leading to larger index sizes and requiring significant resources for graph reconstruction when new alleles are introduced. Here, we propose STR-PG, a topologically decoupled genome-wide framework that separates stable locus representation from scalable STR allele content. STR-PG uses topologically fixed pointer nodes to represent each target locus, while allele sequences, repeat counts, motif annotations, and population frequency metadata are stored in an external registry. Short reads are mapped to STR loci via syncmer-based flanking anchors, and genotyping is performed within a locus-specific candidate space using allele-level alignment likelihood and Bayesian inference. Newly supported alleles can be integrated through registry-level updates without the need to rebuild the graph structure. Evaluations using simulated whole-genome sequencing data, 1000 Genomes Project (1kGP) samples, and r real whole-exome sequencing data from matched whole-blood-cell controls demonstrate that STR-PG maintains accurate genotyping results across various STR classes, reproduces expected population structures, and substantially reduces the computational cost of integrating additional alleles. STR-PG provides a compact and scalable framework for population-scale STR analysis using short-read sequencing.

## Introduction

Short tandem repeats (STRs) account for approximately 3% of the human genome and represent one of its most polymorphic classes of genetic variation [1–3]. Variation in STR length and sequence composition contributes to gene regulation, population diversity, and a broad spectrum of human phenotypes and diseases [3,4]. Despite their biological importance, STRs remain difficult to genotype from short-read sequencing data. Repetitive sequence content reduces unique alignment, whereas length-variable alleles are frequently represented incompletely by a single linear reference, leading to reference bias, alignment ambiguity, and systematic errors in repeat-length estimation [5,6]. Dedicated callers, including HipSTR [7], GangSTR [8], ExpansionHunter Denovo [9], and ExpansionHunter [10], address parts of this problem through local realignment, paired-end evidence, or locus-specific sequence analysis. Nevertheless, their analyses remain anchored to predefined linear-reference loci and allele models, which limits the representation of diverse population haplotypes and newly observed STR alleles [11].

Pangenome graphs provide a natural framework for reducing this reference dependence because they represent sequences from multiple haplotypes within a shared graph [12–14]. Generic graph frameworks, such as Minigraph-Cactus, vg, and GraphTyper [15–18], encode genomic variation as alternative paths and thereby allow reads to be evaluated against a broader set of haplotypes. The graph-construction frameworks used for methodological comparison are summarized in Supplementary Section S3. At highly polymorphic STR loci, however, this otherwise effective representation creates a distinct design problem. In conventional path-based graphs, every STR allele is simultaneously an allelic state and a topological object. As additional haplotypes are incorporated, the number of repeat-length paths, cycles, and locally redundant traversals can grow rapidly, producing dense “hairball” structures and complicated reverse-node alignments (Figure 1A–C). Unlike most SNPs and short indels, STR allele sets are high-cardinality, cohort dependent, and potentially open ended. Consequently, explicitly coupling allele enumeration to graph topology can enlarge indexes, increase the search space for read mapping, and require graph reconstruction whenever a previously unrepresented allele is added. The underlying limitation is therefore not merely computational overhead; it is a mismatch between two quantities with different stability: genomic locus connectivity is comparatively stable, whereas the catalog of STR alleles continually expands as new samples and populations are analyzed.

**Figure 1.**
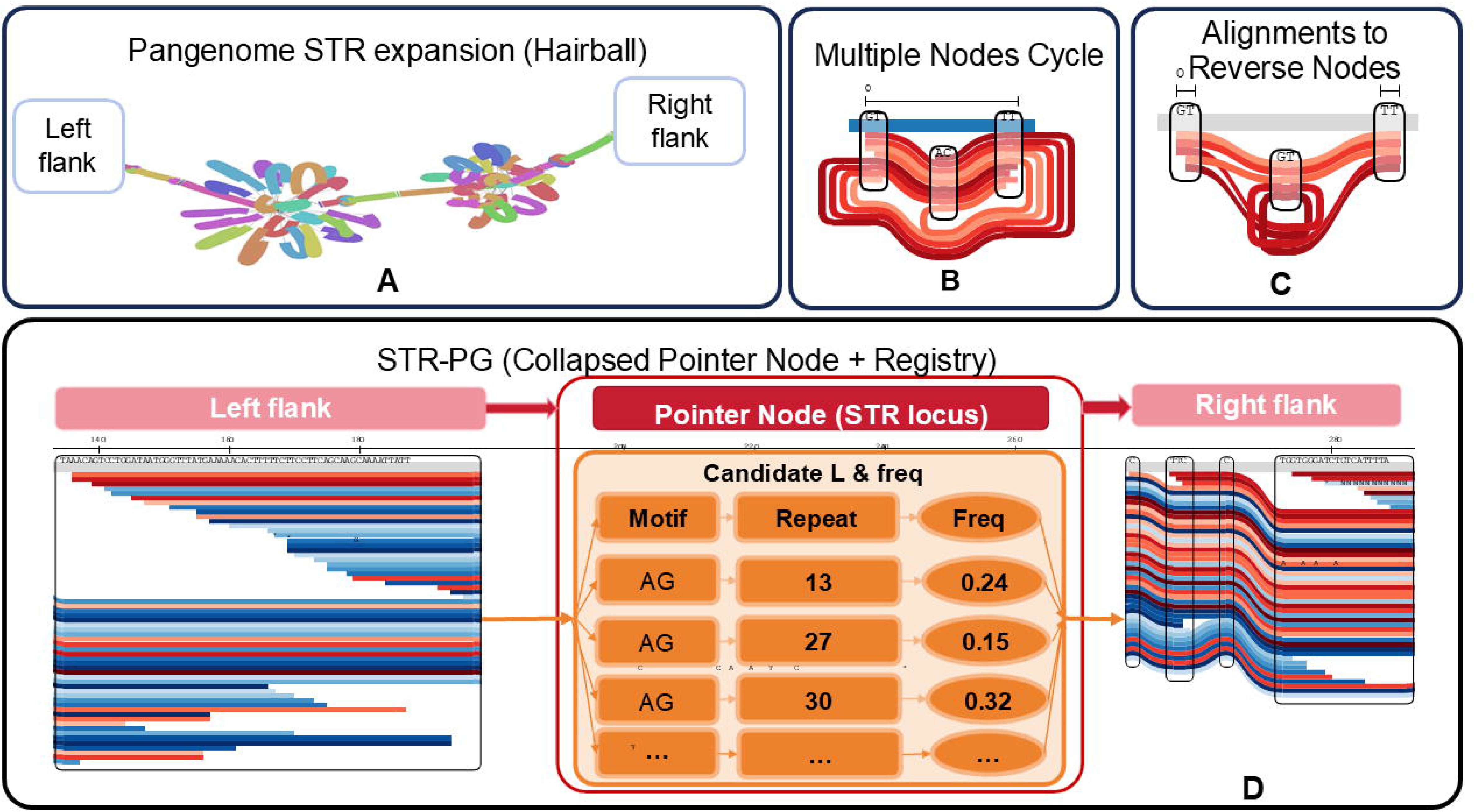
Topology–allele coupling at STR loci and the STR-PG representation. **A**. Explicit representation of numerous STR alleles as alternative paths can produce dense local “hairball” structures in a pangenome graph, visualized using Bandage [33]. **B**. A cyclic STR representation containing multiple repeat-node traversals, visualized using SequenceTubeMap [34]. **C**. Reverse-oriented node traversals can further complicate local read alignment, visualized using SequenceTubeMap [34]. **D**. STR-PG collapses the allele-rich local structure into a fixed-topology pointer node. The pointer preserves connectivity between polymorphic flanking regions, whereas allele sequences, repeat counts, motif annotations, and frequency metadata are stored in an external registry and retrieved for locus-specific genotype inference. **Alt text:** Four-panel schematic comparing conventional STR graph representations with the topology-decoupled STR-PG design. Panel A shows numerous multicolored alternative paths forming dense hairball-like structures between the left and right flanks of a pangenome STR locus. Panel B shows multiple red traversal paths through a cyclic repeat-node representation. Panel C shows alignments involving reverse-oriented nodes, which further complicate local graph traversal. Panel D shows the STR-PG representation, in which the left and right flanks are connected through a single fixed-topology pointer node. An external registry associated with the pointer node stores candidate motif sequences, repeat counts, and allele frequencies, illustrated by AG alleles with repeat counts of 13, 27, and 30 and frequencies of 0.24, 0.15, and 0.32. The design retains flanking connectivity while separating variable STR allele content from graph topology.

Existing strategies do not directly resolve this topology–allele coupling. The danbing-tk toolkit uses repeat-pangenome graphs to improve short-read mapping and characterize VNTR length and motif composition [19]; however, repeat diversity remains encoded within locus-specific graph structures constructed from a predefined set of haplotype-resolved assemblies. PanGenie combines a haplotype-resolved pangenome panel with short-read k-mer counts to reduce the computational burden of mapping-based genotyping [13], but inference is restricted to variants and haplotype paths already represented in the input panel. Long-read tools such as TRGT provide more complete observations of tandem-repeat sequence composition and methylation from PacBio HiFi data [20], yet they are designed around predefined repeat loci and do not by themselves remove the representational cost of encoding each newly observed allele as an additional graph path. Detailed descriptions and parameter settings of the evaluated STR callers are provided in Supplementary Section S2. Thus, these approaches are optimized for repeat-aware mapping, panel-based inference, or long-read characterization, rather than for separating stable graph topology from an independently extensible STR allele catalog. This distinction motivates a more fundamental design question: can a pangenome preserve graph-based locus context while allowing STR allelic diversity to expand independently of graph topology?

To address this problem, we developed STR-PG (Short Tandem Repeat Pangenome Graph), whose central design principle is topology–allele decoupling. Rather than enumerating all STR alleles as alternative graph paths, STR-PG collapses the allele-rich local subgraph into a fixed-topology pointer node. The node preserves the identity of the STR locus, its connectivity to the left and right flanks, and the local haplotypic context (Figure 1D). Allele-resolved information, including nucleotide sequences, repeat counts, motif annotations, and population-frequency metadata, is maintained in an external registry indexed by the pointer node. The graph therefore specifies where the locus is located and how it connects to neighboring sequence, whereas the registry specifies which alleles are currently available for inference. During analysis, short reads are first localized to the corresponding pointer locus using informative flanking anchors. Registry records for that locus are then retrieved to construct a locus-specific candidate allele space for probabilistic genotyping. When the read evidence supports an allele absent from the current catalog, the allele can be incorporated at the registry level without adding another graph path or rebuilding the graph index.

This representation changes how STR diversity scales within a pangenome. Because each STR locus is represented by a single pointer node regardless of the number of observed alleles, local graph topology remains stable as cohort diversity increases. Registry-level updates separate routine allele expansion from costly graph reconstruction, while locus-specific candidate retrieval retains the allele sequence information required for short-read genotyping. In addition, metadata such as motif structure and population frequencies can be extended without modifying the graph backbone, allowing the representation to evolve across diverse populations and successive cohorts. In this study, we investigate whether topology–allele decoupling can alleviate the dense “hairball” structures produced by complex STR variation, reduce their burden on pangenome construction and downstream STR genotyping, and control graph growth and update cost while maintaining accurate STR genotyping in simulated whole-genome data and population-scale human sequencing datasets. STR-PG therefore provides a representation principle for reconciling stable pangenome topology with dynamic, multi-population allelic diversity.

## Results

Complete dataset manifests, software versions, quality thresholds, ablation configurations, and computational environments are reported in Supplementary Section S7 and Tables S4–S7.

### Resolving intra-sample heterogeneity in a multi-bubble simulation

A fundamental limitation of many STR genotyping methods is the strict diploid constraint. Because these methods usually report no more than two alleles per locus, additional allele components caused by intra-sample heterogeneity or sample contamination may be obscured. To evaluate whether STR-PG could resolve such signals, which appear as multiple local bubbles in an allele representation, we designed a controlled simulation of somatic repeat instability.

We generated a synthetic mixture at a (CGC)*n* locus. The sample contained a heterozygous germline genotype of 15/18 repeat units and an expanded subclone spanning 30–40 repeat units (Figure 2A). Standard linear alignment is challenged by these expanded alleles because reads derived from repeat tracts longer than 100 bp often contain insufficient unique flanking sequence for unambiguous placement.

**Figure 2.**
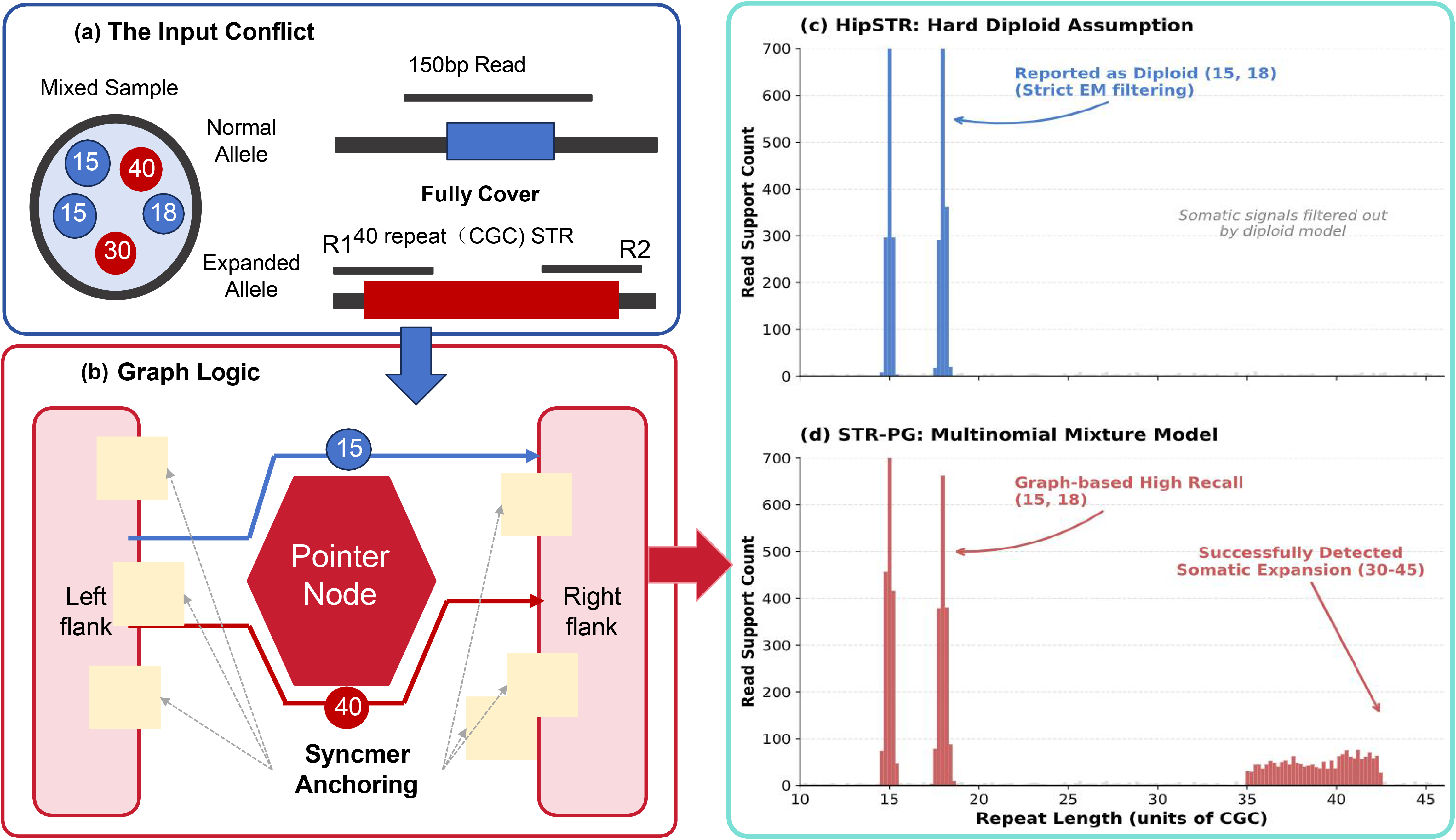
Resolution of intra-sample heterogeneity by STR-PG. **A**. The input conflict: A biological sample with intra-sample heterogeneity contains stable germline alleles (15 and 18 repeat units) together with a heterogeneous population of expanded alleles (30–40 repeat units). Standard 150-bp reads derived from the expanded region often fail to align to a linear reference because they lack sufficient unique flanking context. **B**. Graph logic: STR-PG uses pointer Nodes flanked by unique syncmer anchors. Reads are anchored to the flanking regions, allowing the variable repeat content to be threaded through the pointer Node without requiring a pre-existing path in the graph topology. **C**. Resulting genotype landscape: The output histogram of read support counts shows successful detection of the “multi-bubble” state, including two sharp peaks corresponding to the germline alleles (15 and 18 repeat units) and a broad plateau corresponding to the somatic expansion (30–40 repeat units). **Alt text:** Three-panel schematic illustrating the resolution of intra-sample STR heterogeneity. Panel A shows a mixed sample containing germline alleles of 15 and 18 CGC repeat units and expanded alleles represented by a 40-repeat example. A 150-bp read can fully cover a normal allele but cannot span the complete expanded repeat tract while retaining sufficient flanking sequence. Panel B shows the STR-PG graph logic. Syncmer anchors in the left and right flanks localize reads to a central pointer node, allowing read evidence for both a 15-repeat allele and a 40-repeat allele to be associated with the same STR locus without requiring separate graph paths. Panel C shows a histogram of read-support count against repeat length, with two narrow blue peaks at 15 and 18 repeat units and a broader red distribution spanning approximately 30–40 repeat units.

STR-PG addressed this problem through its pointer-node architecture. Unique syncmers first anchored reads to the conserved left and right flanks (Figure 2B). The internal repeat sequence was then evaluated against locus-specific candidate alleles rather than being forced onto a predefined graph path. Different repeat lengths could therefore be recorded through the pointer node and its associated allele registry.

The read-support distribution contained two sharp peaks at 15 and 18 repeat units, recovering the baseline germline genotype, together with a broad plateau from 30 to 40 repeat units (Figure 2C). The multinomial mixture model identified a broad repeat-length distribution spanning 30–40 repeat units, which was separated from the two major germline allele peaks and retained as an additional heterogeneous signal. Thus, STR-PG retained a simple traversal path in the core graph while representing the complex allele set {15, 18, 30–40} in the external registry. This separation reduced the need to encode extreme allelic diversity directly in graph topology.

### Benchmarking on a high-fidelity simulated cohort

We used GSDcreator [21] to generate a synthetic benchmark that approximated realistic whole-genome sequencing conditions. The dataset had 30× coverage and included 1,200 independent sample–locus records with diverse motif lengths and repeat counts. Empirical models of PCR stutter and sequencing error were also incorporated.

Genotype concordance was stratified by allele length (Figure 3A). For STRs shorter than the read length, STR-PG achieved the highest concordance (0.98), followed by HipSTR (0.94), ExpansionHunter (0.86), GangSTR (0.85), and GraphTyper (0.60). For alleles longer than the read length but shorter than the fragment length, ExpansionHunter and STR-PG showed comparable performance, with concordance values of 0.97 and 0.96, respectively, exceeding GangSTR (0.88), HipSTR (0.65), and GraphTyper (0.60). For expansions longer than the fragment length, ExpansionHunter achieved the highest concordance (0.72), followed by STR-PG (0.64) and GangSTR (0.53); results were unavailable for HipSTR and GraphTyper.

**Figure 3.**
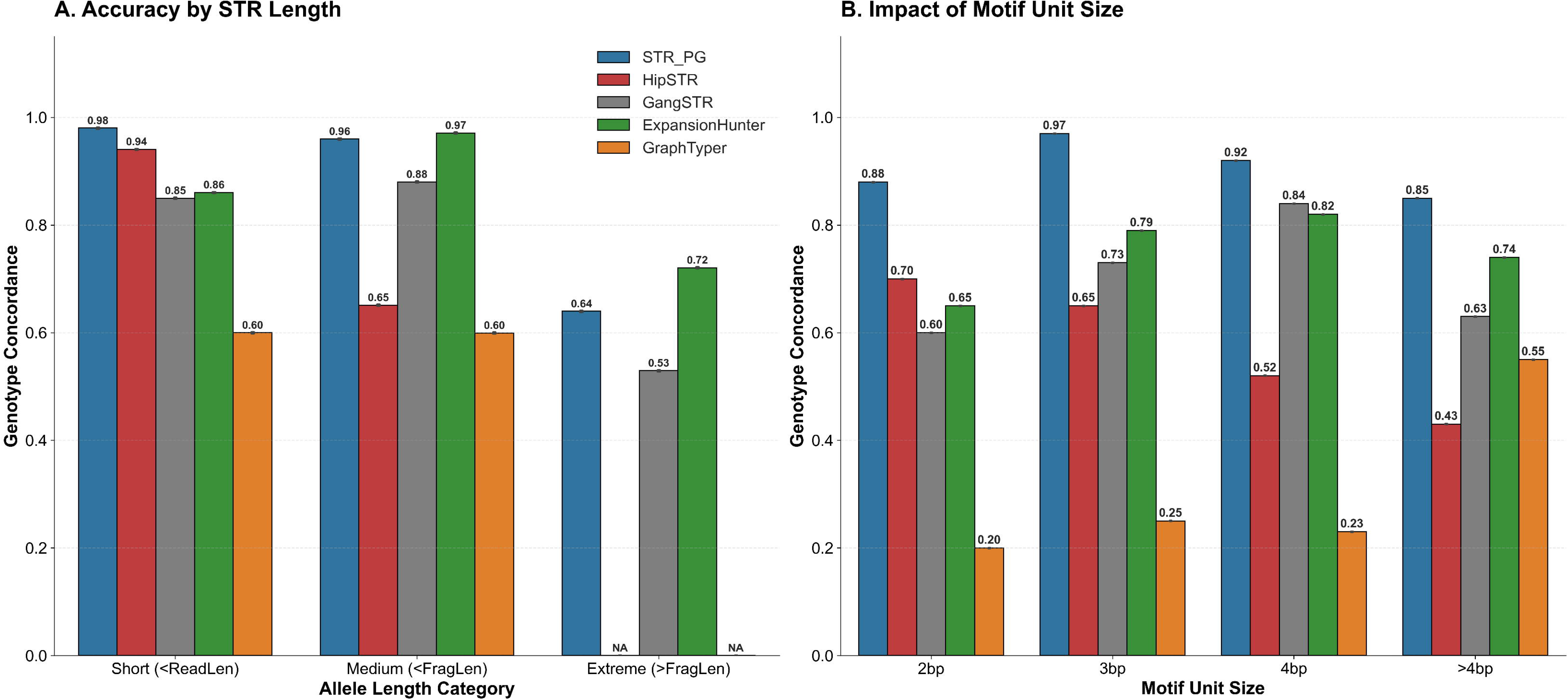
Benchmarking STR genotyping accuracy on simulated whole-genome sequencing (WGS) data. **A.** Genotype concordance stratified by allele length: short (< read length), intermediate (< fragment length), and extreme (> fragment length). **B**. Impact of motif unit size (2–6 bp) on genotyping accuracy. **Alt text:** Two grouped bar charts compare genotype concordance among STR-PG, HipSTR, GangSTR, ExpansionHunter, and GraphTyper on simulated whole-genome sequencing data. Panel A stratifies performance by allele length. For short alleles, concordance values are 0.98, 0.94, 0.85, 0.86, and 0.60, respectively. For medium-length alleles, values are 0.96, 0.65, 0.88, 0.97, and 0.60. For expansions longer than the fragment length, ExpansionHunter has the highest concordance at 0.72, followed by STR-PG at 0.64 and GangSTR at 0.53; HipSTR and GraphTyper are marked as unavailable. Panel B stratifies concordance by motif unit size. STR-PG records 0.88, 0.97, 0.92, and 0.85 for 2-bp, 3-bp, 4-bp, and greater-than-4-bp motifs, respectively, and remains among the highest-performing methods across all motif categories.

We next examined performance across motif lengths (Figure 3B). Dinucleotide repeats were the most difficult class for all tools, consistent with their greater susceptibility to polymerase slippage. Concordance decreased to 0.70 for HipSTR and 0.60 for GangSTR, whereas STR-PG retained a concordance of approximately 0.88. Performance improved for trinucleotide and longer motifs, and STR-PG remained stable across these categories. Overall, the simulation showed that STR-PG performed well for short and intermediate alleles and retained useful sensitivity for expansions beyond the fragment length.

### Evaluation on real WGS data from the 1000 Genomes Project

To assess STR-PG under realistic sequencing noise and population diversity, we analyzed 50 samples from the 1000 Genomes Project (1kGP) [22]. The analysis focused on chromosome 19, which has a high density of STR loci. The cohort represented five superpopulations—AFR, AMR, EAS, EUR, and SAS—and therefore provided a stringent test of traversal across population-diverse haplotypes (Figure 4A). The aggregate number of genotyped STR records was similar among the superpopulations, at approximately 1.71 million, whereas AFR samples showed the expected accumulation of population-specific alleles (503,358). STR-PG indexed and genotyped these variable regions without rebuilding the core graph for each newly observed allele. We compared STR-PG genotypes with the EnsembleTR [23] consensus across genetic backgrounds. Pearson correlations exceeded 0.90 in all representative samples and reached 0.959 for HG00232 (Figure 5A–E). Principal component analysis based only on STR-PG genotypes also recovered the expected population structure. PC1 separated African ancestry, whereas PC2 further resolved Eurasian substructure (Figure 5F). These results indicate that representing STR loci as topological pointers preserved population-informative variation.

**Figure 4.**
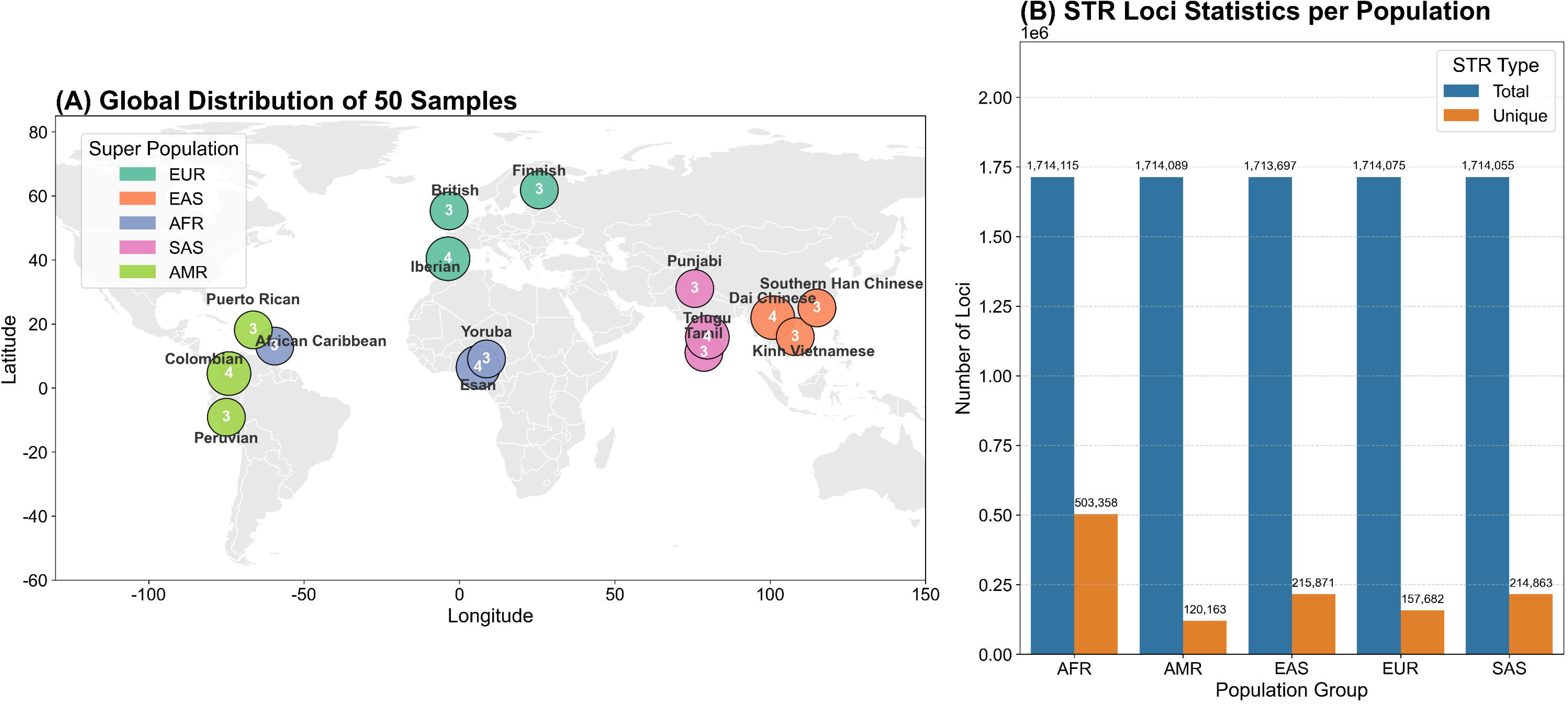
Dataset diversity and locus statistics. **A.** Geographical distribution of the 50 selected 1000 Genomes Project (1kGP) samples, colored by super-population (EUR, EAS, AFR, SAS, and AMR). Bubble size indicates the number of samples in each sub-population. **B**. Number of STR loci genotyped by STR-PG in each super-population. **Alt text:** Two-panel summary of the geographic diversity and STR statistics of 50 samples from the 1000 Genomes Project. Panel A is a world map showing sampled subpopulations in Europe, East Asia, Africa, South Asia, and the Americas. Bubbles are colored by the five superpopulations EUR, EAS, AFR, SAS, and AMR, and the number inside each bubble indicates the number of samples from that subpopulation. Labeled groups include British, Finnish, Iberian, Yoruba, Esan, African Caribbean, Punjabi, Telugu, Tamil, Dai Chinese, Southern Han Chinese, Kinh Vietnamese, Puerto Rican, Colombian, and Peruvian populations. Panel B presents paired bars labeled Total and Unique for each superpopulation. Total values are approximately 1.714 million in all five groups, whereas Unique values are 503,358 for AFR, 120,163 for AMR, 215,871 for EAS, 157,682 for EUR, and 214,863 for SAS.

**Figure 5.**
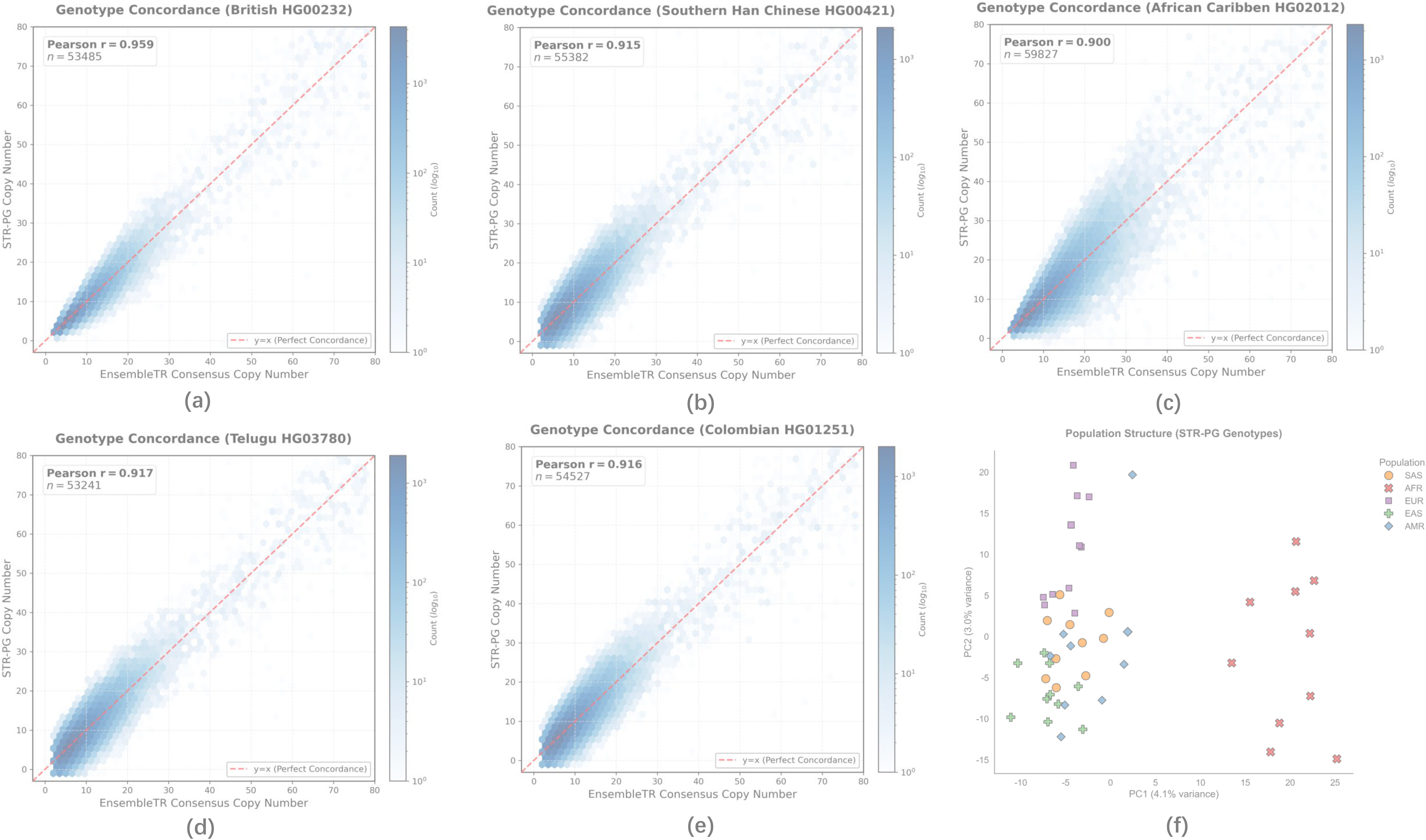
Genotype concordance and population structure. Only loci satisfying the harmonized callable-region, locus-matching, and quality criteria were included. **A-E.** Pairwise scatter plots of STR copy numbers (STR-PG versus the EnsembleTR consensus) for five representative samples from the chromosome 19 analysis of the full 50-sample cohort. The samples shown are: **A.** EUR (HG00232), **B.** EAS (HG00421), **C.** AFR (HG02012), **D.** SAS (HG03780), and **E.** AMR (HG01251). Color intensity indicates locus density on a logarithmic scale. **F.** PCA plot derived from STR-PG genotypes, showing population stratification. **Alt text:** Six-panel comparison of STR-PG genotypes with EnsembleTR consensus genotypes across five representative 1000 Genomes Project samples, followed by population-level principal component analysis. Panels A–E are density scatter plots of EnsembleTR consensus copy number on the horizontal axis and STR-PG copy number on the vertical axis. A red dashed diagonal indicates perfect concordance, and darker blue regions indicate higher observation density. Pearson correlations are 0.959 for British sample HG00232 with 53,485 loci, 0.915 for Southern Han Chinese sample HG00421 with 55,382 loci, 0.900 for African Caribbean sample HG02012 with 59,827 loci, 0.917 for Telugu sample HG03780 with 53,241 loci, and 0.916 for Colombian sample HG01251 with 54,527 loci. Panel F shows a PCA plot of STR-PG genotypes, with samples colored and shaped by superpopulation. African samples are primarily separated along PC1, while the remaining populations show additional structure along PC2.

STR-PG also detected loci with broad, high-entropy read-support profiles that resembled the multi-bubble pattern observed in the simulation. Such heavy-tailed distributions may arise from somatic expansion, cell-line artifacts, or other sources of non-diploid signal. Their biological interpretation therefore requires orthogonal validation. Nevertheless, the results show that STR-PG can retain and quantify complex local evidence that would otherwise be difficult to summarize under a strict diploid model. Dataset versions, registry resources, and locus-harmonization procedures are described in Supplementary Section S5.

### Cross-tool concordance in WES data from matched WBC controls

To evaluate genotyping stability in clinically generated short-read data, we analyzed 50 whole-exome sequencing (WES) datasets derived from whole-blood-cell (WBC) samples used as matched normal controls for patients with solid tumors [24]. The datasets were provided by Geneseeq and were analyzed in a de-identified form. Only WBC-derived sequencing reads were used; tumor sequencing data and clinical phenotypes were not included in this study. Because WES coverage is restricted by the capture design, the comparison was limited to chromosome 19 and chromosome 22 STR loci that satisfied the shared callable-region and coverage criteria defined in Supplementary Section S7 and Tables S5.

STR-PG was benchmarked against HipSTR [7], GangSTR [8], and ExpansionHunter [10] after harmonizing genomic coordinates, repeat motifs, and diploid genotype representations, consistent with established cross-caller comparison practices [11,25]. Under strict locus matching, STR-PG showed the strongest overall agreement with HipSTR. Among 36,408 comparable genotypes, exact genotype concordance was 81.5%, the Pearson correlation was 0.991, and the mean absolute error was 0.159 repeat units (Figure 6A). Exact concordance with GangSTR and ExpansionHunter was 67.2% and 46.7%, respectively, with corresponding mean absolute errors of 0.801 and 3.045 repeat units (Figure 6B,C).

**Figure 6.**
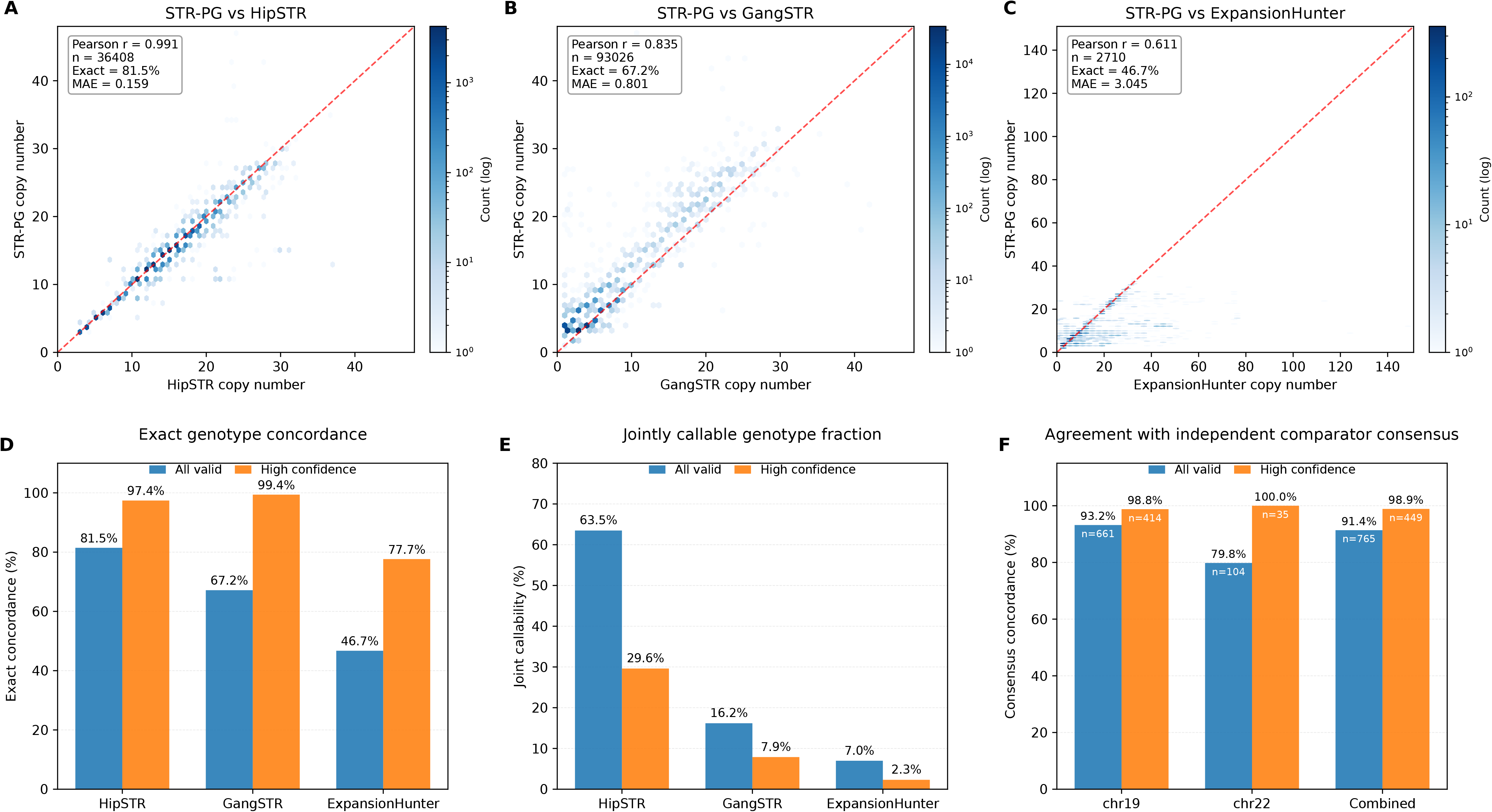
Pairwise concordance between STR-PG and three independent STR callers in WBC sequencing data. **A–C.** Hexbin plots comparing the longer-allele copy number reported by STR-PG and HipSTR, GangSTR, or ExpansionHunter at exactly matched loci. The dashed line indicates identity. Pearson correlation was calculated using longer-allele copy numbers, whereas exact concordance and genotype mean absolute error were calculated from complete diploid genotypes. n denotes the number of sample–locus records with valid genotype measurements from both callers. **D.** Exact diploid genotype concordance among jointly called records. **E.** Joint callability, defined separately for each comparator as the proportion of eligible sample–locus pairs at exactly matched loci for which both STR-PG and the corresponding comparator produced valid genotypes. The denominator was comparator-specific because the number of loci exactly matched to the STR-PG target set differed among callers. High-confidence analysis required both genotype calls to pass the predefined confidence criteria. **F.** Agreement between STR-PG and the consensus genotype formed by the independent comparator callers. n indicates the number of records for which a comparator consensus was available. Comparator consensus was used as complementary evidence and was not considered a ground-truth genotype set. **Alt text:** Six-panel analysis of cross-tool STR genotype concordance in 50 matched-control WBC whole-exome sequencing samples. Panels A–C are density scatter plots comparing STR-PG copy numbers with HipSTR, GangSTR, and ExpansionHunter, with red dashed identity lines. HipSTR shows Pearson r of 0.991, exact concordance of 81.5%, and mean absolute error of 0.159 across 36,408 calls. GangSTR shows r of 0.835, concordance of 67.2%, and error of 0.801 across 93,026 calls. ExpansionHunter shows r of 0.611, concordance of 46.7%, and error of 3.045 across 2,710 calls. Panel D shows that high-confidence filtering raises concordance to 97.4%, 99.4%, and 77.7%, respectively. Panel E shows all-valid and high-confidence jointly callable fractions of 63.5% and 29.6% for HipSTR, 16.2% and 7.9% for GangSTR, and 7.0% and 2.3% for ExpansionHunter. Panel F shows STR-PG agreement with an independent comparator consensus, reaching 91.4% for all valid calls and 98.9% for high-confidence calls in the combined chromosome 19 and 22 analysis.

Quality filtering substantially increased cross-tool concordance. Among high-confidence calls, exact concordance reached 99.4% for GangSTR, 97.4% for HipSTR, and 77.7% for ExpansionHunter (Figure 6D). HipSTR retained the largest shared callable set, whereas the high-confidence call sets from GangSTR and ExpansionHunter covered fewer loci (Figure 6E). An independent majority consensus was constructed from genotypes supported by at least two of the three comparator tools. STR-PG agreed with this consensus for 91.4% of all valid calls and 98.9% of high-confidence calls (Figure 6F). The remaining discrepancies were concentrated mainly at low-confidence, low-depth, or tool-dependent loci.

### Systematic ablation analysis of STR-PG modules

To evaluate the effects of different STR-PG inference components and configurations, we compared eight predefined configurations across 1,200 sample–locus records. The complete SW-based model, S0, served as the reference configuration. S1, S2, and S6 were composite configurations that modified multiple prior-related settings, whereas S3–S5 provided controlled comparisons of the Balding–Nichols (BN) correction, repeat-length smoothing, and likelihood backend, respectively. S7 differed from S0 only by disabling the post-inference mixture diagnostic (Figure 7A). S0 achieved an exact diploid genotype concordance of 92.33% (1,108/1,200; Figure 7B).

**Figure 7.**
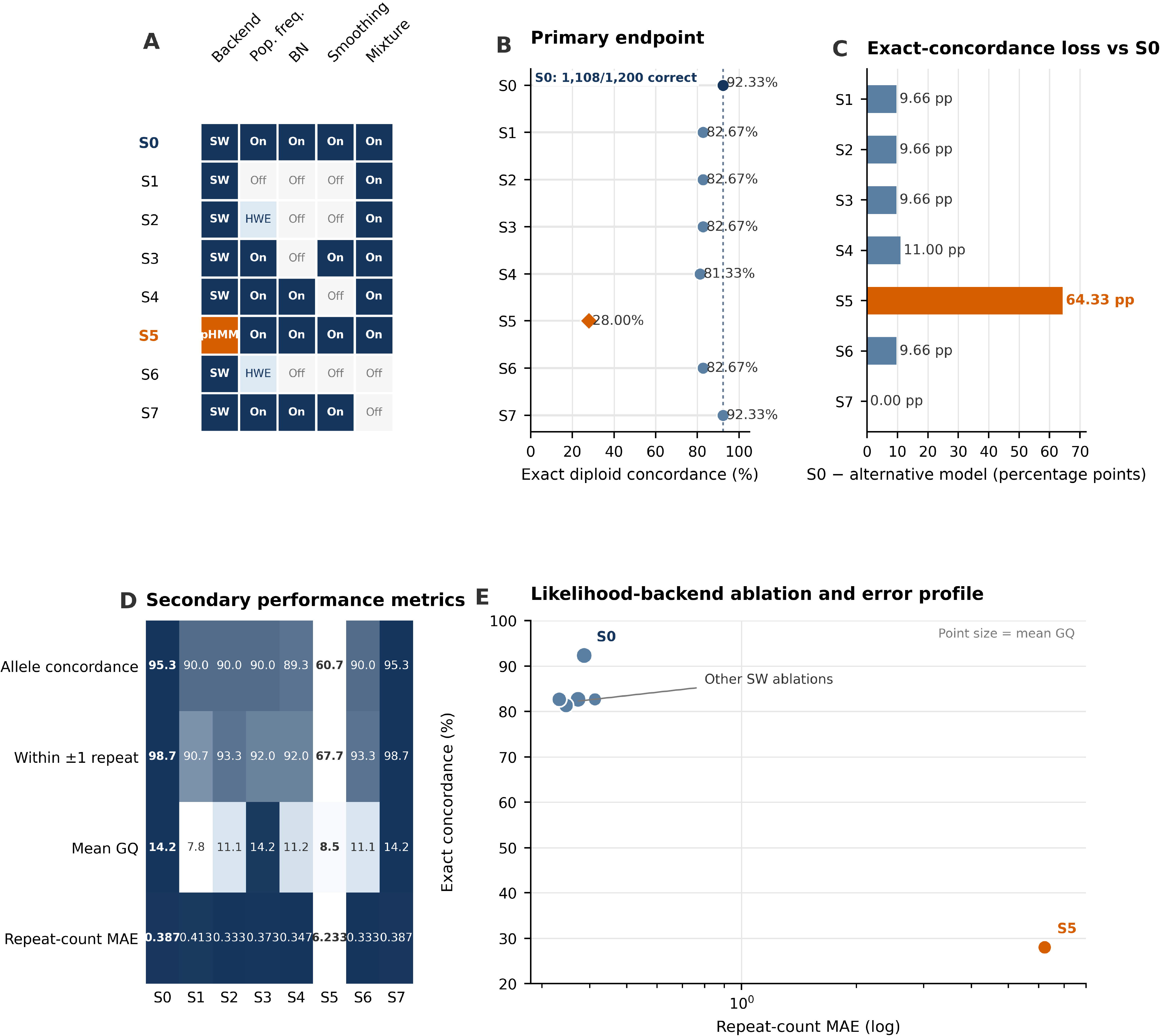
Systematic ablation analysis of STR-PG. **A.** Configurations S0–S7, with S0 representing the complete model. **B.** Exact diploid genotype concordance across 1,200 sample–locus records. **C.** Concordance loss relative to S0. **D.** Secondary performance metrics, including allele concordance, ±1-repeat concordance, mean GQ, and repeat-count MAE. **E.** Relationship between repeat-count MAE and exact concordance. Replacing the SW backend with pHMM in S5 caused the largest performance decline. **Alt text:** Five-panel systematic ablation analysis of STR-PG using 1,200 sample–locus records. Panel A is a configuration matrix for models S0–S7. S0 uses the Smith–Waterman backend with population frequency, Balding–Nichols correction, repeat-length smoothing, and mixture modeling enabled. Models S1–S4 and S6–S7 remove or modify one or more auxiliary components, whereas S5 replaces Smith–Waterman with the experimental pHMM backend. Panel B shows exact diploid concordance of 92.33% for S0, 81.33%–82.67% for the other Smith–Waterman configurations, and 28.00% for S5. Panel C shows concordance losses of approximately 9.66–11.00 percentage points for the Smith–Waterman ablations and 64.33 points for S5. Panel D is a heatmap of allele concordance, within-one-repeat concordance, mean genotype quality, and repeat-count mean absolute error. S0 has 95.3% allele concordance and an error of 0.387, whereas S5 has 60.7% allele concordance and an error of 6.233. Panel E plots exact concordance against log-scaled repeat-count error; S0 occupies the high-concordance, low-error region, while S5 is separated at low concordance and high error.

Among the SW-based configurations, disabling BN correction in S3 reduced exact concordance to 82.67%, corresponding to a decrease of 9.66 percentage points relative to S0. Disabling repeat-length smoothing in S4 reduced exact concordance to 81.33%, a decrease of 11.00 percentage points. The composite configurations S1, S2, and S6 each achieved an exact concordance of 82.67%; however, because these configurations modified multiple prior-related settings simultaneously, their performance differences could not be attributed to a single component. In contrast, S7 produced the same exact concordance as S0, with no change in allele concordance, within-one-repeat concordance, mean GQ, or repeat-count MAE. The identical results for S0 versus S7, and for S2 versus S6, confirmed that the post-inference mixture diagnostic did not affect diploid genotype calling (Figure 7B–D).

The likelihood backend produced the largest performance difference among the evaluated one-factor changes. Replacing the default Smith–Waterman (SW) backend [26] with the candidate pHMM backend in S5 [27] reduced exact concordance from 92.33% to 28.00%, a decrease of 64.33 percentage points (Figure 7C). Allele concordance declined from 95.33% to 60.67%, within-one-repeat concordance declined from 98.67% to 67.67%, and repeat-count MAE increased from 0.387 to 6.233 (Figure 7D). In the error–concordance analysis, S0 and the other SW-based configurations were located in the region of relatively low repeat-count error and high exact concordance, whereas S5 was clearly displaced toward higher error and lower concordance (Figure 7E). These results identify the SW likelihood backend as the largest performance determinant among the evaluated one-factor changes in the current implementation. BN correction and repeat-length smoothing also improved exact concordance in their corresponding controlled comparisons, whereas the mixture diagnostic served only as a post-inference characterization module and did not improve diploid genotype accuracy.

### Performance and scalability

We evaluated the scalability of STR-PG at two distinct levels. First, a controlled regional benchmark was conducted on a 1-Mb region of chromosome 19 containing the highly variable STR loci CACNA1A and GIPC1. Across 112 samples, STR-PG completed initial graph and registry construction in 2.30 s with 48.1 MB peak RAM. Registering one additional sample required 1.14 s and 56.5 MB peak RAM. These measurements were used for the cross-tool comparison with vg and minigraph and are reported in Supplementary Section S1 and Table S1.

Second, we recorded resource consumption during an operational whole-genome workflow constructed from GRCh38 and 1000 Genomes Project resources. Initial construction of the pointer-augmented graph required 21.36 h, whereas registry-level incorporation of one additional sample required 4.3 s and did not require reconstruction of the core graph. The resulting reusable graph index occupied 6.3 GB. Processing one 30× whole-genome sample generated a 114.23-GB GAF-compatible read-localization file because detailed read-level localization and diagnostic fields were retained. These whole-genome measurements were obtained under a different execution setting from the regional cross-tool benchmark and are therefore reported separately in Supplementary Section S7 and Table S7.

Together, these results indicate that STR-PG shifts routine allele integration from graph reconstruction to a lightweight registry-level operation. However, storage of detailed read-localization records remains a practical consideration for population-scale analysis.

### Topological simplification and storage characteristics

The pointer-node architecture reduced the local topological expansion caused by highly polymorphic STR loci. By collapsing complex STR subgraphs into linear logical entities, STR-PG kept local graph complexity independent of the number of registered alleles. The GFA index for the 1kGP cohort occupied 6.3 GB, demonstrating substantial compression of the core graph representation.

Processing one 30× whole-genome sample generated a 114.23-GB GAF-compatible read-localization file because detailed pointer-level localization and diagnostic fields were retained. Thus, STR-PG reduced the size and complexity of the reusable graph index, although storage of full alignment output remained a practical consideration for large cohorts.

## Discussion

Pangenome representation at repetitive loci involves an inherent trade-off. Explicit encoding of population diversity often creates computationally intractable hairball-like graph structures. STR-PG addresses this problem through a pointer-node architecture that decouples graph topology from allele content. This design provides a more stable representation of highly polymorphic repeat regions than explicit path enumeration. It also allows new alleles to be added through registry updates without repeated reconstruction of the core graph.

STR-PG also reduces the sensitivity loss associated with linear-reference alignment. Existing STR callers may be affected by reference bias when the flanking sequence differs from the linear reference genome. STR-PG combines a pangenome framework with syncmer-based seeding. This strategy maintains stable read anchoring even when the flanking regions contain SNPs or small insertions and deletions. Such pangenome-guided anchoring reduces the effect of local sequence divergence on genotyping accuracy. It may therefore recover STR alleles that would otherwise be lost because of alignment clipping or ambiguous placement.

The ablation analysis further showed that STR-PG performance depends on the joint contribution of graph representation, candidate allele construction, and statistical correction. Smith–Waterman-based candidate likelihood evaluation was particularly important for accurate repeat-length estimation. In contrast, the population prior, Balding–Nichols correction [28], length smoothing, and mixture model mainly improved candidate discrimination and genotyping stability.

Beyond conventional diploid genotyping, STR-PG preserves broad or multimodal read-support distributions. This feature provides a basis for detecting intra-sample heterogeneity and may be useful for studying somatic mosaicism [29], repeat expansions, and microsatellite instability. However, non-diploid signals may also arise from PCR stutter, alignment ambiguity, sample contamination, or cell-line artifacts. STR-PG should therefore be considered a framework for detecting and quantifying complex local signals rather than a standalone method for biological or clinical interpretation. Confirmation with paired samples or long-read sequencing remains necessary.

STR-PG is still limited by the properties of short-read sequencing. For expansions that extend well beyond the fragment length, reads cannot span the entire repeat region. Repeat-length estimation may therefore depend on indirect evidence. Syncmer-based anchoring [30,31] also requires sufficiently unique flanking sequences and may become ambiguous in segmental duplications or other low-complexity regions. In addition, although pointer nodes substantially reduce the complexity of the core graph index, GAF-compatible read-localization files that retain detailed pointer-level evidence and diagnostic fields can still require considerable storage.

Overall, STR-PG shows that separating stable locus topology from dynamically expanding allele content can balance compact pangenome representation, incremental update capability, and short-read STR genotyping. Future work should integrate long-read evidence, improve the probabilistic likelihood backend, and extend the intra-sample heterogeneity model to the precise detection of microsatellite instability in cancer genomes. These developments may further improve the analysis of complex repeat regions and somatic STR variation.

## Materials and methods

We introduce STR-PG, a hybrid pangenome architecture formally represented as *G*^′^ = (*G*_core_, *R*). The framework is designed to address the representational complexity and read-alignment ambiguity that can arise when highly polymorphic short tandem repeat (STR) alleles are encoded directly in standard pangenome graphs [5,6,13,14,19,20]. As illustrated in Figure 8, the workflow comprises four modular stages: (a) graph construction and topological decoupling; (b) indexing and alignment; (c) probabilistic genotyping; (d) dynamic registry updating. These stages are followed by output reporting of the maximum-a-posteriori genotype, genotype quality, and supporting diagnostic statistics. Detailed definitions of the mathematical notation and implementation-specific procedures, including the population-aware genotype prior, anchoring threshold, seed chaining, pointer-level locus assignment, Smith–Waterman and pHMM likelihood backends, mixture-model diagnostic, GT and GQ calculation, and registry-update criteria, are provided in Supplementary Section S6 and Table S3.

**Figure 8.**
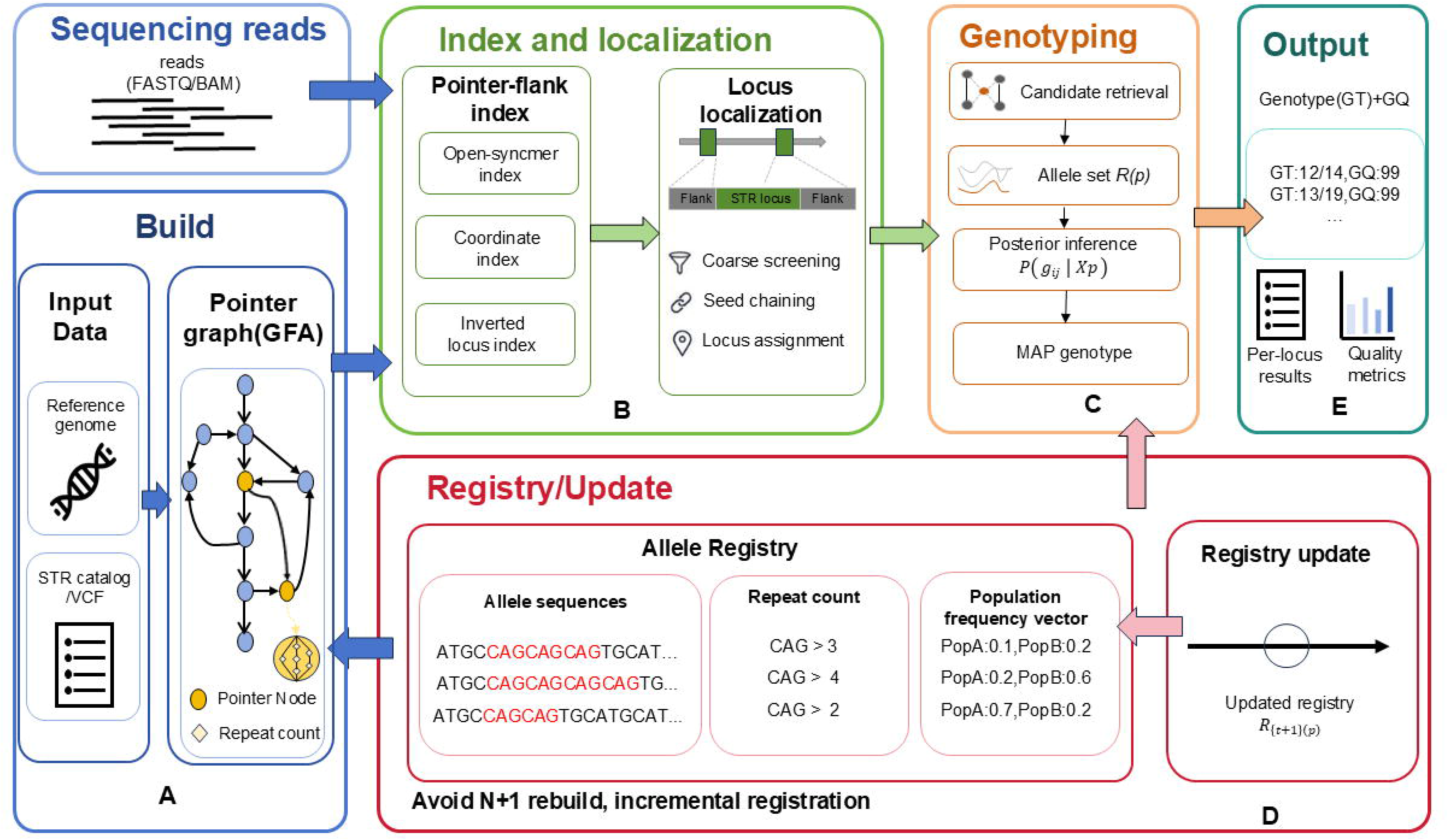
End-to-end workflow of the STR-PG framework. The workflow comprises four core computational stages followed by output reporting. **A.** Build: Hypervariable STR loci are topologically decoupled from the core pangenome graph and represented as fixed-topology pointer nodes. **B.** Index/Map: Syncmer-based seeding and chaining assign sequencing reads to candidate pointer loci and generate GAF-compatible pointer-level locus-localization records. **C.** Genotype: Registry-derived candidate alleles are evaluated using the default Smith–Waterman likelihood backend and population-aware Bayesian inference. **D.** Registry/Update: Allele sequences, repeat counts, motif annotations, population-frequency metadata, and support counts are maintained in an external registry that can be updated without reconstructing the core graph. **E.** Output: STR-PG reports the maximum-a-posteriori genotype, posterior-derived genotype quality, and optional diagnostic statistics. **Alt text:** End-to-end STR-PG workflow organized into five labeled components connected by directional arrows. Panel A shows graph construction from a reference genome and STR catalog or VCF. STR loci are represented as pointer nodes within a GFA graph rather than as multiple allele-specific paths. Panel B shows indexing and localization of FASTQ or BAM sequencing reads using an open-syncmer index, coordinate index, and inverted locus index, followed by coarse screening, seed chaining, and pointer-locus assignment. Panel C shows genotyping through candidate retrieval from the allele registry, construction of a locus-specific allele set, posterior inference, and selection of the maximum-a-posteriori genotype. Panel D shows the external registry, which stores allele sequences, repeat counts, population-frequency vectors, and updated registry records. Registry updates occur incrementally without rebuilding the core graph. Panel E shows output fields including diploid genotype, genotype quality, per-locus results, and quality metrics.

### Graph Construction and Topological Decoupling

In a standard pangenome graph *G* = (*V*, *E*), the high polymorphism of STRs can produce severe topological defects. For a given STR locus *L*, let the allele set be *A* = {*a*_1_, *a*_2_, …, *a_n_*}. As shown in Figure 1A-C, existing graph-construction strategies encounter two major mathematical and computational limitations when encoding *A*.

Redundancy and static representation of the hairball structure (Figure 1A). Explicitly encoding each observed STR allele or population haplotype as an independent graph path increases the number of allele-associated paths and metadata records as the represented cohort expands [13,14,19]. Moreover, a graph constructed from a fixed reference panel does not automatically contain alleles absent from that panel, including newly observed or sample-specific STR alleles.

Alignment ambiguity in cyclic-repeat representations (Figure 1B,C). Representing repeat units as cyclic graph structures permits multiple possible traversals through the same repeat region. As the number of allowable repeat traversals increases, the alignment search space and the ambiguity of determining a unique alignment endpoint can also increase [14,19].

To address these limitations, we define a topological simplification operator *ψ* that maps a highly variable STR subgraph to a single logical entity, as shown in Figures 1D and 8A:

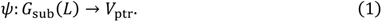

Here, *V*_ptr_ denotes a pointer node. As illustrated in Figure 1D, the STR subgraph is topologically collapsed into a single linear entity connecting the left and right flanks, thereby reducing the local topological complexity to a constant level.

The STR sequence content is decoupled from the graph topology and stored in a dynamic allele registry, R, as shown in Figures 1D and 8D. For each STR locus L, the registry entry indexed by the corresponding pointer-node identifier is defined as:

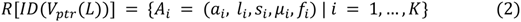

As shown in Figure 8D, the registry explicitly stores three types of information: (1) allele sequences; (2) repeat-count mappings; and (3) population-frequency vectors.

This architecture supports a map-once strategy: reads are anchored to *V*_ptr_ only once, after which complex repeat structures are resolved through registry queries, avoiding redundant realignment against multiple repeat representations.

### Syncmer-based Indexing and Anchoring

To reduce the effect of local variation in STR flanking sequences on read alignment, we use the indexing strategy shown in Figure 8B. The graph index comprises three core components: syncmer seeds, a coordinate index, and a locus-level inverted index.

For an STR locus L, let *S_f_lank*(*L*) denote the indexed syncmer set extracted from the flanking anchor sequences, and let *S_r_* denote the syncmer set extracted from read *r*. After duplicate seed hits are removed and chain consistency is evaluated, let *H*(*r*, *L*) denote the set of unique informative syncmer hits supporting locus *L*. The normalized anchoring score, locus-specific threshold, and corresponding acceptance rule are defined as:

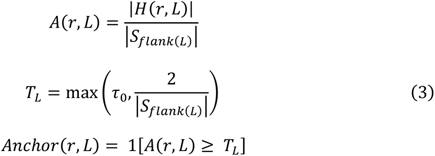

Here, *τ*_0_ is the global minimum anchoring-support fraction applied consistently across all loci and datasets. The term 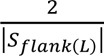 ensures that an accepted anchoring hypothesis contains at least two informative, chain-consistent flank-syncmer hits. Although the numerical value of *T_L_* may vary with the number of indexed flank syncmers available at locus L, the same predefined calculation rule is applied genome-wide and is not optimized separately for individual loci or datasets.

This procedure links coarse syncmer-based seeding to GAF-compatible pointer-level locus localization. Candidate pointer loci are subsequently evaluated using the chaining and pointer-competition procedures described in Supplementary Sections S6.3 and S6.4. The resulting localization record identifies the most strongly supported pointer locus and its mapping confidence; it does not represent a base-resolved alignment to a complete candidate allele path.

### Motif-aware Probabilistic Genotyping

As shown in Figure 8C, the genotyping module receives locus-anchored reads and a registry-derived set of candidate alleles. It then uses a Bayesian framework to infer the maximum a posteriori genotype, following the general probabilistic formulation used by modern STR genotyping methods [7,8,10].

#### Population-aware Genotype Prior

The genotype prior is derived from the population-frequency metadata stored in the allele registry. Population-specific allele frequencies were initialized from webSTR and population genomic resources from the 1000 Genomes Project [2,22]. For an unordered diploid genotype *G_ij_* = {*a_i_*, *a_j_*}, let *p_i_* and *p_j_* denote the normalized frequencies of alleles *a_i_* and *a_j_*, respectively, in the selected population. The baseline Hardy–Weinberg prior is defined as:

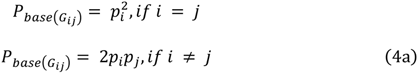

Equation 4a represents only the baseline allele-frequency component. In the complete STR-PG configuration, this baseline is adjusted using a Balding–Nichols correction to reduce overconfidence arising from population differentiation. A repeat-length similarity term is additionally included to account for the tendency of STR alleles to differ through stepwise gains or losses of repeat units. The final normalized genotype prior is defined as:

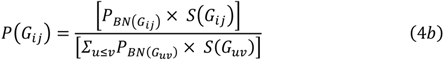

Here, *P_BN_*_(*G*_*_ij_*_)_ denotes the Balding–Nichols-corrected genotype-frequency term, and *S*(*G_ij_*) denotes the repeat-length similarity term. The denominator sums over all unordered diploid genotypes constructed from the registry-defined candidate allele set. Complete definitions of *P_BN_*_(*G*_*_ij_*_)_, *S*(*G_ij_*), and their parameters are provided in Supplementary Section S6.1. The normalized prior is combined with the genotype likelihood to calculate the posterior probability of each candidate diploid genotype.

#### Likelihood Calculation with Default SW and Optional pHMM

STR-PG primarily calculates the likelihood ℒ(ℛ ∣ *GT*) using local Smith-Waterman (SW) alignment (Figure 8C) [26]. For each read *r* and candidate allele ℎ, STR-PG aligns the read to the corresponding allele-specific sequence template and obtains the optimal local alignment score:

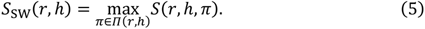

Here, *Π*(*r*, ℎ) denotes the set of possible local alignment paths between read *r* and candidate allele template ℎ, and *S*(*r*, ℎ, *π*) is the score of path *π*, determined by the match, mismatch, gap-opening, and gap-extension scores.

The SW alignment scores are then transformed into normalized read-level likelihoods over the candidate allele set:

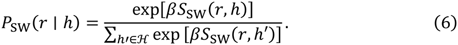

Here, ℋ denotes the candidate allele set at the STR locus, and *β* is a score-scaling parameter controlling the conversion of alignment scores into relative probabilities. The resulting values are normalized compatibility weights derived from relative alignment scores rather than fully calibrated read-generation probabilities.

STR-PG also retains a pair hidden Markov model as an optional candidate-allele evaluation backend. Hidden Markov models provide a probabilistic framework for marginalizing over alternative sequence-alignment paths [27]. When enabled, the pHMM includes match, insertion, and deletion states and can incorporate parameters representing sequencing errors and repeat-associated stutter or slippage, which are important sources of uncertainty in short-read STR analysis [7,8,10,27].

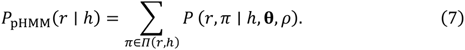

Here, **θ** denotes the transition and emission parameters of the pHMM, and *ρ* is the repeat-slippage or stutter parameter.

Under the default SW model, the allele-specific likelihood used by STR-PG is:

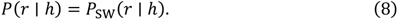

When the optional pHMM is enabled, the allele-specific likelihood is instead defined as:

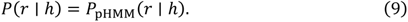

For a diploid genotype *GT* = (ℎ_1_, ℎ_2_), the likelihood contributions of the two alleles are combined using a balanced diploid model:

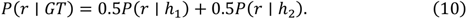

For a homozygous genotype, ℎ_1_ and ℎ_2_ represent the same candidate allele. Assuming conditional independence among reads at a locus given the genotype, the genotype likelihood is calculated as:

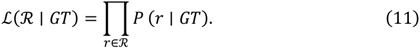

In the current implementation of STR-PG, SW is the default likelihood engine because it provides stable genotyping performance across STR loci at relatively low computational cost. The pHMM is retained as an optional candidate model for evaluating repeat-associated alignment uncertainty and is not required in the default STR-PG workflow.

#### intra-sample heterogeneity Modeling

Read evidence at some STR loci cannot be adequately represented by a conventional diploid model. Such broadened or multimodal repeat-length distributions may reflect somatic mosaicism, but may also arise from amplification stutter, sequencing error, or read-mapping ambiguity [7,8,29]. To retain evidence for these potentially heterogeneous local signals, STR-PG fits a multinomial mixture model over the candidate allele set.

The mixture model uses the same read-level allele likelihood matrix generated during diploid genotyping. Allele contribution weights are estimated using an expectation–maximization procedure, allowing multiple repeat-length components to be represented within a single STR locus. The output of this module includes the estimated allele mixture proportions, supported repeat-length components, and a heterogeneity score summarizing deviation from the diploid expectation.

This module is designed as a diagnostic extension rather than an alternative genotype caller. The reported genotype field (GT), posterior probability, and genotype quality (GQ) are derived from the diploid Bayesian model and are not modified by the mixture analysis. Therefore, the mixture output provides additional information for identifying complex local repeat patterns while maintaining compatibility with conventional VCF-based diploid genotype reporting [32] (Figure 8E).

### Dynamic Registry Updating

To address the N+1 scalability bottleneck, STR-PG does not rebuild the core pangenome graph, represented by the blue component in Figure 8A. As indicated by “Avoid N+1 rebuild” in Figure 8D, when a new allele record *A_new_* is accepted, it is added only to the registry entry associated with the corresponding pointer locus. The accepted allele record is defined as:

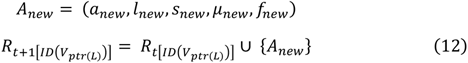

This registry-level operation avoids reconstruction of the core graph. Its computational cost depends on the number of accepted allele records and the corresponding registry operations rather than on rebuilding the graph topology.

### Scalable Implementation

To exploit the constant-size topological representation provided by the pointer-node architecture, STR-PG is implemented in Python using memory-efficient data structures.

Hash-based external allele registry. The allele registry is central to the scalability of STR-PG. We use canonical sequence storage to reduce memory consumption. Rather than storing repeated instances of identical allele strings for each sample, which requires *O*(*N* × *L*) space, identical allele sequences are assigned unique identifiers in a global string pool, and only (AlleleID, Frequency) pairs are stored. This reduces the approximate memory complexity to *O*(*U* × *L*), where *U* is the number of distinct alleles in the population. Because *U* ≪ *N* at most loci, this strategy substantially reduces memory usage.

Memory-efficient syncmer index. During graph alignment, STR-PG uses open syncmers instead of conventional minimizers. Because syncmers are selected according to their positions within a window, they are more robust to mutations near sequence boundaries. To improve tolerance to SNPs and insertions or deletions in STR flanking sequences, we implement a redundant-seed strategy. STR-PG does not depend on a single anchor; instead, it extracts a set of candidate syncmers from each flanking window. Anchoring is determined by voting and requires only a subset of seeds to map to the core graph, thereby reducing the effect of local sequence differences.

### Graph visualization

Graph structures were inspected and visualized using Bandage [33] and SequenceTubeMap [34].

## Authors’ contributions

JW conceived and designed the model. JY and ZX implemented the code, performed the experiments, and curated the data. HT participated in the sequence alignment. All authors read and approved the final manuscript.

## Data availability

The public 1000 Genomes Project and EnsembleTR resources analyzed in this study are described in References [22,23]. The WBC WES datasets contain controlled human genomic information and cannot be deposited in a public repository because of participant-privacy, institutional, and data-use restrictions. Access to these data may be considered upon reasonable request to Jiajing Yuan, subject to approval by the data-providing institution, the relevant ethics committee, and an appropriate data-use agreement. The simulated datasets, locus manifests, processed benchmark tables, and analysis scripts used to reproduce the reported figures and summary statistics are available through the STR-PG software repository and its associated archived release.

## Code availability

The STR-PG source code, test data, configuration files, and analysis scripts are available at https://github.com/YJ124/STR_PG. The analyses reported in this study were performed using STR-PG version 1.0.0

## Acknowledgments

This work was supported in part by funds from the National Natural Science Foundation of China (NSFC: 62572389 and T2541083).

## Supplementary material

### Supplementary File 1 STR-PG supplementary materials

File name: STR-PG_Supplementary_Materials.docx; Microsoft Word format. This file contains Supplementary Sections S1–S7, Supplementary Tables S1–S7, and Supplementary References.

**Table S1 Quantitative assessment of graph construction and update performance**

**Table S2 Conceptual positioning of STR-PG relative to representative methods**

**Table S3 Symbols and notation used in the main text and Supplementary Section S6**

**Table S4 The 50 1000 Genomes Project samples used in the population-scale evaluation**

**Table S5 Cross-tool call definitions, quality thresholds, and genotype-normalization rules**

**Table S6 STR-PG ablation configurations**

**Table S7 Operational whole-genome resource measurements**

