## Supplementary Materials for "STR-PG: A Topology-decoupled Pangenome Framework for Scalable Short-read Genotyping of Short Tandem Repeats"

#### Supplementary S1. Performance Benchmarking

##### 1. Comparative Analysis of Construction Efficiency

A fundamental challenge in population-scale pangenomics is the "N+1" scalability bottleneck—the requirement to re-align or re-index the entire graph whenever novel samples are integrated. To evaluate how STR-PG addresses this limitation, we conducted a performance benchmark using a 1,000,000 bp region of Chromosome 19 (GRCh38), specifically targeting the high-entropy STR loci *CACNA1A* and *GIPC1*. This benchmark is regional rather than genome-wide and is intended to provide a controlled cross-tool comparison for graph construction and update behavior.

Our benchmarking reveals that while general-purpose tools like Minigraph [S1] are highly efficient for structural variants, they encounter a substantial "computational wall" when resolving small, repetitive STR alleles across 112 samples. Minigraph required 1112.094 seconds to complete the graph induction. This latency is primarily driven by the necessity of VCF-to-FASTA reconstruction and iterative sequence-to-graph alignment. In contrast, STR-PG directly parses variant records into a decoupled registry, completing the end-to-end construction in 2.30 seconds. This 483-fold speedup underscores that bypassing the alignment phase is essential for maintaining tractability in repetitive genomic regions.

##### 2. Topological Complexity and Memory Footprint

Memory management remains a critical metric for the accessibility of pangenome workflows. Traditional frameworks such as vg [S2] explicitly encode every observed allele as a unique path, which frequently leads to exponential topology explosions (the "hairball" problem) in highly polymorphic regions.

As illustrated by our data, vg's peak memory usage surged during the GFA conversion phase to 108.4 MB for a single megabase. When scaled to 112 samples with Minigraph, memory consumption peaked at 1208.3 MB. STR-PG, however, utilizes fixed Pointer Nodes to represent STR loci, offloading sequence diversity to an external registry. This architectural design maintains a minimal memory footprint of 48.1 MB, effectively decoupling the physical graph complexity from the number of registered alleles.

##### 3. Scalability in Incremental Update Scenarios

To simulate real-world growth in clinical or population databases, we measured the cost of adding a single novel sample to an established graph. For vg and Minigraph, even an incremental update necessitates a significant re-calculation of graph topology or indexing buffers, leading to memory spikes of 237.5 MB and 464.8 MB, respectively. STR-PG's Registry-Based Update mechanism circumvents this overhead by performing a metadata-level registration. Integrating a new sample took only 1.14 seconds with negligible memory increase (56.5 MB). These findings suggest that STR-PG's resource requirements remained low in the evaluated incremental-update benchmark, whereas traditional frameworks remain bound by resource demands that scale linearly or higher with the number of samples.

**Table S1 Quantitative assessment of graph construction and update performance.**

| Task | Toolkit | Runtime (s) | Peak RAM (MB) |
|---|---|---|---|
| Initial Construction | vg (v1.60.0) | 2.78 | 108.4 |
| | minigraph (v0.20) | 1112.09 | 1208.3 |
| | STR-PG | 2.30 | 48.1 |
| Incremental Update | vg(v1.60.0) | 8.37 | 237.5 |
| | minigraph(v0.20) | 10.81 | 464.8 |
| | STR-PG | 1.14 | 56.5 |

### 4. Computational Environment

All benchmarks were executed on an Intel(R) Xeon(R) Gold 6248R node (3.00GHz, 512 GB RAM) running Linux (CentOS 7). Resource usage was captured using /usr/bin/time -v to ensure precise measurement of Peak Resident Set Size (RSS).

### Supplementary S2. The descriptions and parameter settings of STR detection tools

We compared STR-PG to representative baselines chosen according to task scope. HipSTR [S3], GangSTR [S4], and ExpansionHunter [S5] were used as short-read STR genotyping baselines; GraphTyper [S6,S7] was included as a graph-centric genotyping baseline; vg and minigraph were used for graph construction and update comparisons in the regional scalability benchmark. Pan-Genie [S8] and prancSTR [S9] are discussed as related methods but were not included as primary end-to-end baselines in the reported experiments because their assumptions and roles differ from those of STR-PG.

*HipSTR* [S3] utilizes a profile hidden Markov model (HMM) [S13] to characterize STR variation by performing haplotype-based realignment. This approach explicitly models PCR stutter and sequencing slippage errors, ensuring high-precision genotyping for alleles that remain within the physical constraints of individual reads. In this study, HipSTR was deployed with its standard parameter set, utilizing pre-aligned BAM files and a reference-integrated STR catalog to generate genotype calls.

*GangSTR* [S4] adopts a comprehensive statistical framework that incorporates multiple sequence signals, including spanning reads and flanking read pairs, into a maximum-likelihood estimation model. By leveraging the insert size distribution of paired-end libraries, it can estimate allele lengths that significantly exceed the fragment length. We operated GangSTR under its default settings to evaluate its robustness in genome-wide STR profiling across varying repeat unit sizes.

*ExpansionHunter* [S5] is specifically engineered to resolve large, often pathogenic, repeat expansions that typically elude standard alignment-based methodologies. It combines a targeted graph-based alignment strategy with specialized logic to resolve repeat counts from "in-repeat" reads and mate-pair information. For the performance evaluation, ExpansionHunter was executed using its default parameters and the manufacturer-provided locus specification files for the targeted STR regions.

*GraphTyper* [S6,S7] introduces a population-scale pangenomic perspective by realigning sequencing reads to localized graph structures containing known variants. Although GraphTyper and GraphTyper2 are general-purpose graph-based variant genotypers rather than dedicated STR callers, they provide graph-centric references for evaluating localized variant representation and population-scale genotyping.

*Pan-Genie* [S8] is a k-mer-based pan-genomic genotyping tool that is particularly adept at inferring various types of variants using pan-genomic reference sequences derived from haplotype resolution. Pan-Genie leverages a haplotype-resolved pangenome reference to infer genotypes by integrating k-mer counts from short-read sequencing data with known haplotype structures. This method significantly mitigates reference bias and enhances the resolution of complex variants, including large insertions and variants located in repetitive genomic regions. For STR-PG analysis, Pan-Genie provides a robust framework to accurately characterize polymorphic loci by exploiting existing population-scale haplotype information.

*prancSTR* [S9] operates downstream of germline TR genotypers (HipSTR) and employs a statistical framework to distinguish genuine mosaic alleles from sequencing-induced stutter artifacts. Notably, this approach enables the detection of mosaicism in individual whole-genome sequencing datasets without requiring matched control samples.

### Supplementary S3. Overview of Pangenome Comparison Frameworks

To contextualize the performance of STR-PG, we benchmarked it against two representative pangenome graph toolkits that reflect different algorithmic approaches to graph construction and variation representation.

#### 1. The vg Toolkit (Variation Graph Toolkit)

*vg* [S2] represents a foundational framework for base-level accurate variation graphs. Its core architecture relies on bi-directed graphs where genomic variations (from SNPs to small indels) are explicitly encoded as alternate paths relative to a reference backbone. This path-centric model ensures high resolution for fine-mapping but is inherently susceptible to "topology explosions" in high-entropy regions, where the combinatorial growth of diverse alleles leads to a "hairball" structure. In our benchmarking, we employed the standard vg construct and vg view workflow using default parameters. This configuration was intended to evaluate how traditional explicit path encoding handles the memory and time requirements of multi-allelic STR loci in a population cohort.

#### 2. Minigraph

*minigraph* [S1] is a computationally efficient tool designed for the induction of pangenome graphs focused on large-scale structural variations (SVs). Unlike base-level tools, minigraph utilizes a minimizer-based heuristic to iteratively align new sequences to an existing graph, adding only those segments that represent significant structural divergence. While this approach provides exceptional scalability for whole-genome comparisons, its default heuristic (typically filtering variants shorter than 50 bp) is not optimized for granular STR genotyping. For this study, minigraph was executed using the -xgg induction workflow under default settings. This serves to demonstrate the baseline performance of structural-centric pangenome builders when confronted with the high-frequency, small-scale insertions and deletions typical of microsatellite evolution.

**Supplementary S4. Conceptual positioning of STR-PG relative to representative methods**

**Table S2 Conceptual positioning of STR-PG relative to representative methods**

| Method Class | Representative Tools | Allelic Representation | Novel Allele Integration | Heterogeneity Modeling |
|---|---|---|---|---|
| Generic pangenome graph toolkits | Minigraph-Cactus[S1]; Vg[S2]; GraphTyper [S6,S7]; | Alleles represented explicitly as graph paths | Requires graph reconstruction or re-indexing | Not explicitly modeled |
| Short-read STR callers | HipSTR[S3]; GangSTR[S4]; ExpansionHunter [S5]; | Locus-specific models anchored to linear reference | Limited to predefined loci or catalogs | Typically assumes diploid genotypes |
| Alignment-free population genotyping | Pan-Genie[S8] | k-mer or haplotype-based inference without explicit graph traversal | Limited support for dynamic allele discovery | Not explicitly modeled |
| Mosaic-aware STR methods | prancSTR [S9] | Linear-reference-based locus models | Not designed for dynamic allele integration | Explicit modeling of non-diploid signals |

Note: This comparison is intended to clarify methodological scope rather than imply universal superiority across all tasks. Several existing methods are optimized for specific applications, such as repeat-expansion detection, population-scale variant genotyping, or downstream mosaic-signal analysis.

### Supplementary S5: Benchmark Datasets and Reference Resources

#### 1. The EnsembleTR Consensus Dataset

To evaluate the performance of Short Tandem Repeat (STR) genotyping, we utilized EnsembleTR [S10] as the primary benchmark consensus dataset. EnsembleTR represents a high-confidence callset generated through the integration of multiple state-of-the-art STR genotyping algorithms, including ExpansionHunter [S5], HipSTR [S3], GangSTR [S4].

EnsembleTR provides an externally generated consensus callset that reduces dependence on any single caller. In this study it is used as the benchmark truth source for sensitivity, precision, and genotype-concordance analyses; its residual uncertainty and the exact release and filtering criteria will be reported with the experimental manifest rather than describing it as an error-free gold standard.

#### 2. Population-frequency resources and registry preparation

The initial STR allele registry used in STR-PG was constructed from the webSTR(http://webstr.ucsd.edu/) resource [S11], which provides population-scale short tandem repeat (STR) variation data derived from whole-genome sequencing across diverse human populations.

For each STR locus, webSTR reports allele-level information including repeat counts, allele sequences (when available), and population-specific allele frequency estimates. In addition, locus-level summary statistics such as heterozygosity and allele count distributions are provided. These data collectively enable initialization of both the candidate allele space and the associated population-aware priors used in STR-PG.

We extracted STR loci and allele records from webSTR using the publicly available dataset corresponding to the GRCh38 reference assembly. For each locus, the following fields were retained:

- Locus identifier and genomic coordinates
- Repeat motif and motif length
- Allele repeat counts
- Population-specific allele frequencies across super-populations
- (When available) allele sequences or repeat representations

To ensure consistency with the STR-PG framework, loci with ambiguous motif definitions or incomplete coordinate information were excluded. In addition, alleles with extremely low support (e.g., observed in only a single individual) may be optionally filtered or down-weighted during initialization to reduce noise in the candidate set. All loci were mapped to STR-PG pointer nodes based on genomic coordinates and motif identity.

For each STR locus L, the extracted allele records were used to initialize the registry entry:

$$R[ID(V_{ptr}(L))] = \{A_i = (a_i, l_i, s_i, \mu_i, f_i) \mid i = 1, \ldots, K\} \quad \text{(S1)}$$

where:

- $a_i$: allele identifier derived from webSTR
- $l_i$: repeat count
- $s_i$: allele sequence or motif-expanded representation
- $\mu_i$: motif annotation, including the repeat unit and motif structure
- $f_i$: population-frequency vector across super-populations

When explicit allele sequences are unavailable, the preparation workflow may reconstruct a motif-expanded candidate from the motif and repeat count together with the locus flanks. Population frequencies are normalized within each population before registry serialization.

The resulting registry defines the candidate allele order used by both likelihood backends. Frequency metadata are optional: they contribute to the genotype prior only when the population-frequency component is enabled.

STR-PG can extend this initial catalog only when novel-allele discovery is explicitly enabled. Accepted alleles and provisional observations are handled by the update rules described below; registry initialization is therefore population informed but not immutable.

### Supplementary S6. Additional methodological specifications for read localization, genotype calling, and registry updating

Symbols used in this section and in the main text are summarized in Table S3.

**Table S3 Symbols and notation used in the main text and Supplementary S6**

| Symbol | Definition |
|---|---|
| **Graph and registry representation** | |
| $\boldsymbol{G = (V, E)}$ | Sequence graph with node set V and edge set E. |
| $\boldsymbol{G' = (G_{core}, R)}$ | Hybrid STR-PG representation comprising the graph-resident core backbone $G_{core}$ and the external allele registry R. |
| $\boldsymbol{G_{core}}$ | Graph-resident backbone used by STR-PG to store stable locus topology and flanking context. |
| $\boldsymbol{G_{sub(L)}}$ | Conceptual explicit allele-rich subgraph at STR locus L, used only to contrast conventional path encoding with the pointer-node design. |
| $\boldsymbol{\psi}$ | Conceptual simplification operator that maps $G_{sub(L)}$ to a single pointer node; the implementation directly constructs the pointer representation. |
| $\boldsymbol{V_{ptr(L)}}$ | Pointer node representing STR locus L in the graph. |
| $\boldsymbol{ID[V_{ptr(L)}]}$ | Unique identifier of the pointer node representing locus L. |
| $\boldsymbol{R}$ | External allele registry indexed by pointer-node identifiers. |
| $\boldsymbol{R\left[ID[V_{ptr(L)}]\right]}$ | Registry entry associated with the pointer node of locus L. |
| $\boldsymbol{A_i = (a_i, \ell_i, s_i, \mu_i, f_i)}$ | Registry record for candidate allele $i$, containing its identifier, repeat count, sequence, motif annotation, and population-frequency vector. |
| $\boldsymbol{a_i}$ | Identifier of candidate allele $i$. |
| $\boldsymbol{\ell_i}$ | Repeat count of candidate allele $a_i$. |
| $\boldsymbol{s_i}$ | Nucleotide sequence or motif-expanded representation of candidate allele $a_i$. |
| $\boldsymbol{\mu_i}$ | Motif annotation of candidate allele $a_i$, including the repeat unit and motif structure. |
| $\boldsymbol{f_i}$ | Vector of population-specific frequencies stored for candidate allele $a_i$. |

| Symbol | Definition |
|---|---|
| $\boldsymbol{K}$ | Number of registered candidate alleles at locus L. |
| $\boldsymbol{A_L = \{a_1, \ldots, a_K\}}$ | Registry-defined candidate allele set associated with locus L. |
| **Reads and syncmer-based anchoring** | |
| $\boldsymbol{r}$ | A sequencing read. |
| $\boldsymbol{R_L = \{r_1, \ldots, r_N\}}$ | Set of reads assigned to locus L. |
| $\boldsymbol{N}$ | Number of reads assigned to locus L. |
| $\boldsymbol{S_r}$ | Syncmer set extracted from read r. |
| $\boldsymbol{S_{flank(L)}}$ | Indexed syncmer set extracted from the flanking anchor sequences of locus L. |
| $\boldsymbol{H(r, L)}$ | Set of unique informative syncmer hits supporting locus L after duplicate removal and chain-consistency filtering. |
| $\boldsymbol{A(r, L)}$ | Normalized anchoring score for read $r\,at$ locus L, computed as $\frac{card[H(r,L)]}{card[S_{flank(L)}]}$. |
| $\boldsymbol{T_L}$ | Anchoring threshold for locus L, defined by the global rule $\max\left[\tau_0, \frac{2}{[S_{flank(L)}]}\right]$. |
| $\boldsymbol{\tau_0}$ | Global minimum anchoring-support fraction applied consistently across loci and datasets. |
| $\mathbf{1}(\boldsymbol{condition})$ | Indicator function equal to 1 when the stated condition is satisfied and 0 otherwise. |
| **Seed chaining and pointer-locus assignment** | |
| $\boldsymbol{h}$ | A seed hit considered during chaining. |
| $\boldsymbol{p}$ | Graph path on which seed hit h is observed. |
| $\boldsymbol{pos_{p(h)}}$ | Path coordinate of seed hit h. |
| $\boldsymbol{pos_{r(h)}}$ | Read coordinate of seed hit h. |
| $\boldsymbol{d_h}$ | Diagonal offset of seed hit h, defined as pos_p(h) - pos_r(h). |
| $\boldsymbol{\delta}$ | Slack parameter controlling diagonal consistency between seed hits; the default value is 10 bp. |

| Symbol | Definition |
|---|---|
| $\boldsymbol{H_L}$ | Number of unique chain-consistent seed hits retained for pointer locus L. |
| $\boldsymbol{span_L}$ | Path span covered by the selected seed chain for pointer locus L. |
| $\boldsymbol{dispersion_L}$ | Root-mean-square dispersion of diagonal offsets within the selected chain for pointer locus L. |
| $\boldsymbol{S_L}$ | Single-read chain score for pointer locus L. |
| $\boldsymbol{S_L^{pair}}$ | Paired-end chain score for pointer locus L after combining mate-specific scores and the paired-support bonus. |
| $\boldsymbol{S_1, S_2}$ | Best and second-best pointer-locus scores used for ambiguity filtering and mapping-quality calculation. |
| $\boldsymbol{MAPQ}$ | Mapping-quality score derived from the margin between the best and second-best pointer-locus scores; this is distinct from genotype quality. |
| **Population-aware diploid genotyping** | |
| $\boldsymbol{G_{ij} = \{a_i, a_j\}}$ | Unordered diploid genotype formed from candidate alleles $a_i$ and $a_j$. |
| $\boldsymbol{G_{2(L)}}$ | Set of all unordered diploid genotypes constructed from $A_L$. |
| $\boldsymbol{p_i, p_j}$ | Scalar frequencies of alleles $a_i$ and $a_j$ in the selected population; these are entries retrieved from $f_i$ and $f_j$. |
| $\boldsymbol{P_{base(G_{ij})}}$ | Baseline Hardy-Weinberg genotype prior for $G_{ij}$. |
| $\boldsymbol{P_{BN(G_{ij})}}$ | Balding-Nichols-corrected genotype-frequency term for $G_{ij}$. |
| $\boldsymbol{\theta}$ | Population-differentiation parameter used in the Balding-Nichols correction. |
| $\boldsymbol{S(G_{ij})}$ | Repeat-length similarity term applied to genotype $G_{ij}$. |
| $\boldsymbol{\sigma}$ | Scale parameter controlling the decay of the repeat-length similarity term with repeat-count difference. |
| $\boldsymbol{P(G_{ij})}$ | Final population-aware genotype prior, normalized over all genotypes in $G_{2(L)}$. |

| Symbol | Definition |
|---|---|
| $P(R_L \mid G_{ij})$ | Likelihood of the read set assigned to locus L under genotype $G_{ij}$. |
| $P(G_{ij} \mid R_L)$ | Posterior probability of genotype $G_{ij}$ given the reads assigned to locus L. |
| **Default Smith-Waterman and optional pHMM likelihood backends** | |
| $T_a$ | Fully expanded sequence template for candidate allele a. |
| $q(r, a)$ | Best Smith-Waterman alignment score for read r against T_a after considering both read orientations. |
| $\ell(r, a)$ | Normalized read-level log compatibility of read r with allele a obtained from the Smith-Waterman score softmax. |
| $F_L, F_R$ | Left and right flanking sequences used to construct an expanded candidate template. |
| $m_a$ | Length of the expanded candidate template T_a. |
| $b_a, e_a$ | Start and end coordinates of the half-open repeat interval in T_a. |
| $\rho_j$ | Region label at template column j, taking the value flank or repeat. |
| $\alpha_\rho, \beta_\rho$ | Region-specific indel-open and indel-extension probabilities for region ρ. |
| $P_X \rightarrow Y^\rho$ | Transition probability from pHMM state X to state Y in region ρ, where X and Y belong to {M, I, D}. |
| $\varepsilon$ | Mismatch probability used in pHMM match-state emissions. |
| $x_i$ | Nucleotide at read position i. |
| $x_j$ | Nucleotide at candidate-template position j. |
| $b_x$ | Background nucleotide probability used for insertion emissions; fixed at 0.25 per nucleotide. |
| $M_i, j, I_i, j, D_i, j$ | Log-space forward variables for the pHMM match, insertion, and deletion states. |
| $n$ | Length of read r. |
| $r^{rc}$ | Reverse complement of read r. |
| $LSE$ | Log-sum-exp operator. |

| Symbol | Definition |
|---|---|
| $logaddexp$ | Numerically stable operation for computing the logarithm of the sum of two exponentials. |
| $\log P_{forward}(r \mid a)$ | Semi-global pHMM forward log likelihood of read r under candidate allele a for one read orientation. |
| $\log P(r \mid a)$ | Orientation-marginalized pHMM log likelihood of read r under candidate allele a. |
| **Optional allele-mixture diagnostic** | |
| $w = (w_1, \ldots, w_K)$ | Non-negative allele-weight vector in the mixture model, with the weights summing to 1. |
| $L_{diploid}^{lik}$ | Best likelihood-only log score under the unordered diploid genotype model. |
| $L_{diploid}^{paper}$ | Best diploid log score after adding the genotype log prior. |
| $L_{mix}$ | Best log likelihood under the flexible K-component allele-mixture model. |
| $D_{likelihood}, D_{paper}$ | Reported mixture diagnostic score differences calculated relative to the likelihood-only and prior-adjusted diploid reference scores, respectively. |
| **Reported genotype fields** | |
| $G_{MAP}$ or $G*$ | Maximum-a-posteriori unordered diploid genotype. |
| $GT$ | Reported genotype field, equal to $G_{MAP}$. |
| $P\_MAP = P(G_{MAP} \mid R_L)$ | Normalized posterior probability of the maximum-a-posteriori genotype. |
| $GQ$ | Posterior-derived genotype quality, calculated from the error probability $1 - P_{MAP}$ and capped at 99. |
| **Novel-allele acceptance and registry updating** | |
| $a_{new}$ | Identifier of a newly accepted allele added to the registry. |
| $\ell_{new}$ | Repeat count of the newly accepted allele. |
| $s_{new}$ | Sequence of the newly accepted allele. |

| Symbol | Definition |
| --- | --- |
| $\boldsymbol{\mu_{new}}$ | Motif annotation of the newly accepted allele. |
| $\boldsymbol{f_{new}}$ | Population-frequency vector assigned to the newly accepted allele. |
| $\boldsymbol{R_t, R_t + 1}$ | Registry states before and after an update step. |
| $\boldsymbol{p_{new}^{P}}$ | Initialized scalar frequency of the newly accepted allele in superpopulation P. |
| $\boldsymbol{\alpha}$ | Pseudo-count smoothing constant used during population-frequency initialization; distinct from the pHMM parameter $\alpha_p$. |
| $\boldsymbol{1(sample\ in\ P)}$ | Indicator of whether the discovery sample belongs to superpopulation P. |
| $\boldsymbol{Count_{P(a)}}$ | Stored support count for allele a in superpopulation P. |
| $\boldsymbol{N_P}$ | Total number of observed allele copies stored for superpopulation P. |

#### S6.1 Population-aware genotype prior

For an unordered diploid genotype $G_{ij} = \{a_i, a_j\}$ constructed from the registry-defined candidate allele set $A_L$, STR-PG defines a population-aware prior using the allele-frequency metadata stored in the allele registry. Let $p_i$ and $p_j$ denote the population-specific frequencies of alleles $a_i$ and $a_j$, respectively. The baseline Hardy–Weinberg prior is defined as:

$$P_{base(G_{ij})} = p_i^2, if\ i = j$$

$$P_{base(G_{ij})} = 2p_i p_j, if\ i \neq j \qquad (S2)$$

To reduce sensitivity to population differentiation, STR-PG applies a Balding–Nichols correction:

$$P_{BN(G_{ij})} = p_i^2 + p_{i(1-p_i)\theta}, if\ i = j$$

$$P_{BN(G_{ij})} = 2p_i p_{j(1-\theta)}, if\ i \neq j \qquad (S3)$$

Here, θ is the population-differentiation parameter. When θ = 0, the Balding–Nichols-corrected prior reduces to the baseline Hardy–Weinberg prior. A positive value of θ increases the prior weight assigned to homozygous genotypes and reduces the assumption that allele frequencies follow an ideal panmictic population. This correction therefore limits overconfidence in equilibrium-based genotype weights when population structure is present.

Because STR alleles commonly differ through gains or losses of repeat units, STR-PG additionally applies a repeat-length similarity term motivated by the stepwise mutation model:

$$S(G_{ij}) = \exp\left[-\frac{(l_i - l_j)^2}{(2\sigma^2)}\right] \quad (S4)$$

Here, $l_i$ and $l_j$ are the repeat counts of alleles $a_i$ and $a_j$, respectively, and σ controls how rapidly the similarity term decreases as the difference in repeat count increases. Genotypes composed of alleles with similar repeat counts therefore receive a larger similarity weight than genotypes composed of alleles with widely separated repeat counts.

The final population-aware genotype prior is obtained by combining the Balding–Nichols-corrected frequency term with the repeat-length similarity term and normalizing the result over all unordered diploid genotypes:

$$P(G_{ij}) = \frac{\left[P_{BN(G_{ij})} \times S(G_{ij})\right]}{\left[\Sigma_{u \le v} P_{BN(G_{uv})} \times S(G_{uv})\right]} \quad (S5)$$

In the denominator, the indices u and v enumerate all candidate alleles in $A_L$, with $u \le v$ ensuring that each unordered diploid genotype is included only once. The resulting prior satisfies:

$$\Sigma_{i \le j} P(G_{ij}) = 1$$

Population-frequency metadata contribute to genotype inference only when the population-prior component is enabled. When this component is disabled, genotype inference is performed without population-frequency weighting. The same registry-defined candidate allele ordering is used for likelihood calculation and prior normalization.

**S6.2 Basis for selecting the anchoring threshold ($T_L$)**

For an STR locus L, let $S_{flank(L)}$ denote the indexed syncmer set extracted from the flanking anchor sequences, and let $S_r$ denote the syncmer set extracted from read r. After duplicate seed hits are removed and chain consistency is evaluated, let $H(r, L)$ denote the set of unique informative syncmer hits supporting locus L. Consistent with Equation 3 in the main text, the normalized anchoring score is defined as:

$$A(r, L) = \frac{|H(r, L)|}{\left|S_{flank(L)}\right|} \quad (S6)$$

where $|H(r, L)|$ is the number of unique, chain-consistent syncmer hits supporting locus L, and $\left|S_{flank(L)}\right|$ is the total number of indexed syncmers extracted from the flanking anchor sequences of that locus.

A read is considered compatible with locus L when:

$$A(r, L) \ge T_L$$

The locus-specific anchoring threshold is defined using a single global rule:

$$T_L = \max\left[\tau_0, \frac{2}{\left|S_{flank(L)}\right|}\right] \qquad (S7)$$

where $\tau_0$ is the global minimum anchoring-support fraction applied consistently across all loci and datasets. The second term, $\frac{2}{|S_{flank(L)}|}$, ensures that an accepted anchoring hypothesis contains at least two informative, chain-consistent flank-syncmer hits.

The threshold is designed to balance two competing requirements. First, it must be sufficiently permissive to tolerate sequencing errors, single-nucleotide variants, and small insertions or deletions in the STR flanking regions. Second, it must be sufficiently stringent to reject off-target matches caused by low-complexity sequences, segmental duplications, or repeated syncmer patterns.

Although the numerical value of $T_L$ may vary with the number of indexed flank syncmers available at locus L, the same predefined calculation rule is applied genome-wide. Therefore, $T_L$ is not independently optimized for individual STR loci or evaluation datasets.

Under the default STR-PG indexing configuration, syncmers are generated using k = 15, s = 5, and t = 2, with 100-bp flanking anchor sequences retained on both sides of each STR locus. A read must provide at least two informative flank-syncmer hits that form a chain-consistent localization hypothesis before it can be promoted to locus-level genotyping.

After the initial compatibility test, the retained seed hits are evaluated using the chaining and pointer-locus competition procedures described in Supplementary Sections S6.3 and S6.4. Reads that do not satisfy the minimum seed-support or chain-consistency requirements are excluded from subsequent genotype likelihood calculation.

**S6.3 Chaining rule for seed consistency**

After coarse seeding, STR-PG groups all seed hits by path and evaluates whether their read coordinates and path coordinates are mutually consistent. For a seed hit $h$ on path $p$, let $p_{\mathrm{pos}}(h)$ and $r_{\mathrm{pos}}(h)$ denote the path and read coordinates, respectively, and define the diagonal offset

$$d(h) = p_{\mathrm{pos}}(h) - r_{\mathrm{pos}}(h) \qquad (S8)$$

Two hits are considered chain-consistent when their diagonal offsets differ by at most a slack parameter $\delta$. In the default mapper this slack is 10 bp.

Operationally, for each path, hits are binned by round$(d(h)/\delta)$. The candidate chain for that path is the densest diagonal bin. The path-level chain is then ordered by increasing path coordinate. Across all paths, the best chain is the one with the largest number of supporting hits; ties are resolved by preferring the chain with larger path span and then smaller within-chain diagonal dispersion. Reads without a chain of at least two informative seeds are not promoted to locus-level genotyping.

This chaining rule formalizes the "Chaining" module in Fig. 8B: it is not merely a speed optimization, but the criterion by which seed matches are converted into a biologically interpretable local alignment hypothesis.

#### S6.4 Pointer-level locus assignment and GAF-compatible output

For each pointer locus, duplicate seed hits are removed within the densest diagonal bin. Let $H_L$ be the number of remaining chain-consistent hits, $\mathrm{span}_L$ the path span covered by those hits, and $\mathrm{dispersion}_L$ the root-mean-square dispersion of their diagonal offsets. The single-read chain score is:

$$S_L = H_L + 0.01\,\mathrm{span}_L - 0.02\,\mathrm{dispersion}_L \qquad (S9)$$

For paired-end data, chains are matched by pointer identifier. If both mates support the same pointer, their scores are summed and a fixed paired-support bonus of 0.5 is added; otherwise, the available mate-specific score is retained.

$$S_L^{\mathrm{pair}} = S_L^{(1)} + S_L^{(2)} + 0.5\,\mathbf{1}\{\text{both mates support } L\} \qquad (S10)$$

Candidate pointer loci are ranked by this score. If a second candidate exists and the difference between the best and second-best scores is below 0.25, the read or read pair is labeled ambiguous and excluded from genotyping.

The mapping-quality field is a deterministic transform of the score margin. With $S^{(1)}$ and $S^{(2)}$ denoting the best and second-best pointer scores,

$$\mathrm{MAPQ} = \min\big(60,\ \mathrm{round}\big[10\max\big(0, S^{(1)} - S^{(2)}\big)\big]\big) \qquad (S11)$$

When no second candidate is present, MAPQ is set to 60. The mapper emits a GAF-compatible locus-localization record containing the query interval, pointer identifier, chain-supported pointer interval, MAPQ, and diagnostic tags for score, seed count, left/right hits, support fraction, dispersion, and mate. This record represents pointer-level localization; it is not a base-resolved alignment to a complete candidate allele path.

This pointer competition is the implemented link between the chaining and genotyping modules in Fig. 8B–C. Allele-specific sequence comparison is deferred to the likelihood backend after locus assignment.

#### S6.5 Definition of the optional expanded-template pHMM backend

The production-default likelihood backend in STR-PG is Smith–Waterman score softmax. A pHMM backend is retained as an explicit experimental option. The implemented pHMM is a linear, allele-specific M/I/D forward model over each fully expanded candidate template; it does not contain a cyclic motif-phase state block.

For candidate allele $a$ with repeat count $l_a$, the template is

$$T_a = F_L \oplus \mathrm{motif}^{l_a} \oplus F_R, \qquad m_a = \mathrm{length}(T_a) \qquad (S12)$$

The candidate stores repeat boundaries $b_a$ and $e_a$, which define a half-open repeat interval. At template column $j$, the region label $\rho(j)$ is "repeat" when $b_a \leq j - 1 < e_a$ and "flank" otherwise. Region-specific indel-open and indel-extension parameters are denoted $\alpha_\rho$ and $\beta_\rho$.

$$P(M \to M) = 1 - 2\alpha_\rho; \quad P(M \to I) = P(M \to D) = \alpha_\rho; \quad P(I \to I) = P(D \to D) = \beta_\rho (S13)$$

The return transitions satisfy $P(I \to M) = P(D \to M) = 1 - \beta_\rho$. The audited defaults are a mismatch rate of $\varepsilon = 0.01$; flank indel-open and indel-extension probabilities of 0.003 and 0.10, respectively; repeat indel-open and indel-extension probabilities of 0.005 and 0.30, respectively; and an insertion background probability of 0.25 per nucleotide.

Match-state emissions use the candidate template base $x_j$:

$$e_M(x_i, x_j) = 1 - \varepsilon \quad (x_i = x_j)$$
$$e_M(x_i, x_j) = \frac{\varepsilon}{3} \quad (x_i \neq x_j) \qquad (S14)$$
$$e_I(x_i) = 0.25$$

Deletion states are silent. The forward variables $M_{i,j}$, $I_{i,j}$, and $D_{i,j}$ are stored as natural-log probabilities. A semi-global uniform start distribution is used across template positions:

$$M_{0,j} = -\log(m_a + 1), \quad j = 0, \ldots, m_a; \qquad I_{0,j} = D_{0,j} = -\infty \qquad (S15)$$

For $i \geq 1$ and $j \geq 1$, letting LSE denote the log-sum-exp operator, the match-state recurrence is

$$M_{i,j} = \log e_M(x_i, x_j) + \mathrm{LSE}(M_{i-1,j-1} + \log P_{MM},\ I_{i-1,j-1} + \log P_{IM},\ D_{i-1,j-1} + \log P_{DM}) (S16)$$

The insertion- and deletion-state recurrences are

$$I_{i,j} = \log 0.25 + \mathrm{LSE}(M_{i-1,j} + \log P_{MI},\ I_{i-1,j} + \log P_{II})$$
$$D_{i,j} = \mathrm{LSE}(M_{i,j-1} + \log P_{MD},\ D_{i,j-1} + \log P_{DD}) \qquad (S17)$$

For read length $n$, semi-global termination sums over all template end positions and all three states:

$$\mathrm{Log}P(r \mid a, \mathrm{forward}) = \mathrm{LSE}_{1 \leq j \leq m_a}(M_{n,j}, I_{n,j}, D_{n,j}) \qquad (S18)$$

Read orientation is marginalized with equal prior weight rather than selected by a maximum:

$$\mathrm{Log}P(r \mid a) = \mathrm{logaddexp}(\log P(r \mid a, \mathrm{forward}),\ \log P(\mathrm{rc}(r) \mid a, \mathrm{forward})) - \log 2 \quad (S19)$$

The start distribution already contains the template-length factor $\frac{1}{(m_a+1)}$; therefore, no additional division by $m_a$ is applied at termination, because doing so would duplicate length normalization and introduce template-length bias. The pHMM returns raw natural-log likelihoods, with larger values indicating greater support. Its mathematical implementation is covered by forward-versus-brute-force, transition-direction, semi-global-boundary, normalization, exact-match-ranking, and no-double-normalization tests. Because current comparative evidence does not establish superiority over the SW backend, pHMM results must be labeled experimental and are not used as the production-default results.

Before likelihood calculation, both backends use the same deterministic informative-read cache. A mapped read is selected only when either orientation connects a 12-bp flank anchor to at least two adjacent motif units with a motif mismatch fraction $\leq 0.10$. If either mate satisfies this boundary criterion, both mates in the pair are selected.

$$q_{r,a} = \max\{\mathrm{SW}(r, T_a), \mathrm{SW}(\mathrm{rc}(r), T_a)\}$$

$$\ell_{r,a} = \mathrm{logsoftmax}_a\left(q_{r,a}/12\right) \qquad (S20)$$

The default SW parameters are match = 2, mismatch = −5, gap = −5, and band = 500. The temperature softmax is normalized across the ordered candidate set for each read. Historical read caps, early stopping after a fixed number of good pairs, candidate pruning, and backend-specific read selection are not part of the current production implementation.

#### S6.6 Optional mixture-model diagnostic for heterogeneous local signals

Some loci may show evidence broader than a diploid model. STR-PG optionally fits a K-component allele mixture as a diagnostic model to summarize heterogeneous repeat-length signals. The diagnostic does not alter the reported diploid GT, posterior, or GQ.

The implementation records two diploid reference scores. The likelihood-only score is

$$L_{\mathrm{diploid}}^{(\mathrm{lik})} = \max_{G \in \mathcal{G}_2} \log P(R_L \mid G) \qquad (S21)$$

and the paper-compatible posterior score is

$$L_{\mathrm{diploid}}^{(\mathrm{paper})} = \max_{G \in \mathcal{G}_2} \left(\log P(R_L \mid G) + \log P(G)\right) \qquad (S22)$$

For non-negative allele weights $\mathbf{w} = (w_1, \ldots, w_K)$ that sum to one, the mixture log likelihood is

$$L_{\mathrm{mix}} = \max_{\mathbf{w}} \sum_{r \in R_L} \log\left(\sum_{k=1}^{K} w_k\, P(r \mid a_k)\right) \qquad (S23)$$

The mixture module reports descriptive statistics, including estimated component proportions and repeat-length distributions. These statistics are used for characterizing heterogeneous loci and are not interpreted as formal hypothesis-testing results. A HET flag is emitted only when a separately calibrated threshold is supplied. No asymptotic chi-square threshold is assumed by default.

#### S6.7 Definitions of GT and GQ

The genotype field (GT) is the maximum-a-posteriori unordered allele pair:

$$\mathrm{GT} = G_{\mathrm{MAP}} = \underset{G \in \mathcal{G}_2}{\operatorname{argmax}}\, P(G \mid R_L) \qquad (S24)$$

The posterior is normalized over every unordered diploid genotype constructed from the registry-defined candidate set.

GQ is computed directly from the posterior error probability of the selected genotype:

$$\mathrm{GQ} = \min\bigl(99, \mathrm{round}\bigl[-10\log_{10}\bigl(1 - P(G_{\mathrm{MAP}} \mid R_L)\bigr)\bigr]\bigr) \qquad (S25)$$

The value is capped at 99. The current implementation does not use a best-versus-second-best score-gap approximation.

Consequently, no separate $\Delta$ statistic is required to reproduce GQ; candidate ordering affects only deterministic tie resolution, not the posterior formula.

GQ summarizes posterior discrimination among candidate diploid genotypes and must not be interpreted as read mapping quality.

Accordingly, Fig. 8E reports the MAP diploid genotype (GT) with its posterior-derived confidence (GQ).

#### S6.8 Acceptance criteria for novel alleles

Novel-allele discovery is disabled by default and must be enabled explicitly. A hypothesis is generated only from a read spanning both 12-bp flank anchors, with repeat-region length divisible by the motif length and motif mismatch fraction $\leq 0.10$. The inferred repeat count must be absent from the registered candidate set and supported by at least three reads.

A temporary candidate is accepted only when all implemented criteria are met: minimum support of three reads; exclusion of a one-repeat-unit stutter shadow (default shadow fraction $\leq 0.15$); log-score gain $\geq 3$ over the registered-only call; GQ $\geq 20$; and inclusion of the new allele in the MAP genotype. Localized hypotheses failing any criterion remain provisional and do not receive learned population frequency.

The distinction between provisional and accepted hypotheses separates exploratory observations from registry learning and prevents unqualified calls from entering the active candidate prior.

#### S6.9 How registry updating avoids error accumulation

Registry updates are designed to accumulate evidence in count space rather than in frequency space. For each population, STR-PG stores allele counts and total observed allele copies and recomputes frequencies from these sufficient statistics whenever the registry is written out. This prevents the repeated rounding and renormalization errors that would occur if previously rounded frequencies were updated recursively.

A second safeguard is confidence filtering. In the default update workflow, only genotypes with GQ above a user-specified threshold are allowed to contribute to the registry, and only homozygous calls are used for direct frequency learning. Homozygous high-confidence loci contribute two allele copies, whereas heterozygous, missing, or low-confidence calls are ignored. This rule trades speed for conservativeness and sharply reduces the risk that an uncertain allele assignment will be amplified by repeated updates.

A third safeguard is pseudo-count smoothing. When historical counts are not available, the initial frequency table is converted to approximate counts using a fixed base total (1000 in the current workflow), after which all subsequent updates are performed on counts. Newly accepted alleles receive only a small initial prior mass until additional samples accumulate. Together, these rules make registry growth incremental while limiting error propagation across update rounds.

### Supplementary S7 Experimental datasets, software settings, and computational environment

#### S7.1 Common reference resources

All analyses used the GRCh38 coordinate system. STR-PG analysis resources were constructed from chromosome-specific GRCh38 FASTA sequences, STR locus catalogs, population allele-frequency tables, and the corresponding pointer-node registry. Unless otherwise stated, each STR locus was represented together with 100-bp flanking sequences on both sides.

The syncmer index used k = 15, s = 5, and t = 2. Repeat motifs were canonicalized across cyclic rotations and reverse-complement representations before cross-tool locus matching or genotype comparison. STR intervals used in the comparison workflow were normalized to 1-based inclusive coordinates.

For the WBC cross-tool benchmark, HipSTR, GangSTR, and ExpansionHunter used a common reference containing GRCh38 chromosomes 19 and 22. The SHA-256 checksum of the combined reference FASTA was:

```
be7970813a9e2d7afcccede5a5682b8d5a8d55c66ab4450e14ad01e4f7fea7f9
```

The original comparator resources were hg38.hipstr_reference.bed.gz for HipSTR, hg38_ver13.bed.gz for GangSTR, and eh_hg38_variant_catalog.json for ExpansionHunter. Chromosome-specific chr19 and chr22 subsets derived from these resources were used in the completed analysis.

#### S7.2 Controlled multi-bubble simulation

The within-sample heterogeneity experiment simulated a (CGC)n STR locus containing two major alleles of 15 and 18 repeat units and an expanded component spanning 30-40 repeat units. Simulated 150-bp paired-end short reads were used to evaluate the ability of STR-PG to preserve and distinguish multimodal repeat-length evidence at a single locus.

The experiment was designed to represent local read evidence that departs from a conventional diploid assumption. Reads generated from the different repeat-length components produced multiple separable peaks in the local repeat-length spectrum. STR-PG aggregated the pointer-localized evidence and used the optional mixture model to characterize the two major alleles and the expanded component.

The primary outputs were the retention of the simulated repeat-length components, separation of the read-length modes, and identification of a heterogeneous repeat-length distribution by the mixture diagnostic.

#### S7.3 Simulated cohort benchmark

GSDcreator was used to generate a 30× simulated short-read cohort containing 1200 independent STR sample-locus records [S14]. The benchmark covered repeat motifs of 2-6 bp and included alleles shorter than the read length, alleles between the read and fragment lengths, and alleles longer than the fragment length.

All evaluated configurations used the same simulated read evidence, candidate allele set, genotype enumeration procedure, and truth labels. Genotypes were treated as unordered diploid allele pairs, and phase was ignored.

The primary endpoint was exact diploid genotype concordance with the simulated truth. Additional metrics included allele repeat-count deviation, mean absolute error, genotype quality, and performance stratified by motif length and allele-length class.

#### S7.4 1000 Genomes Project evaluation

The population-scale evaluation included 50 publicly available 1000 Genomes Project samples, with 10 samples selected from each of the AFR, AMR, EAS, EUR, and SAS superpopulations [S15]. The cohort covered 15 populations: ACB, ESN, and YRI in AFR; CLM, PEL, and PUR in AMR; CDX, CHS, and KHV in EAS; FIN, GBR, and IBS in EUR; and ITU, PJL, and STU in SAS. Analysis was restricted to chromosome 19.

GRCh38-aligned CRAM files were obtained from the European Nucleotide Archive using the run accessions listed in Table S4. Population and superpopulation assignments followed the official 1000 Genomes Project Phase 3 sample panel. The geographic descriptions in Table S4 refer to the population definition or sample-collection location used by the International Genome Sample Resource and do not represent individual-level birthplace information.

STR-PG genotypes were compared with the EnsembleTR consensus call set used in the population analysis [S16]. Before comparison, chromosome names, genomic intervals, canonical repeat motifs, and repeat-count representations were harmonized. The two alleles of each diploid genotype were sorted before comparison, and phase was ignored. Matched loci were used to calculate genotype concordance, repeat-count deviation, and Pearson correlation. STR-PG genotypes were also used for population principal component analysis.

**Table S4 The 50 1000 Genomes Project samples used in the population-scale evaluation**

| Sample ID | ENA run accession | Population code | Population/region | Superpopulatio n |
|---|---|---|---|---|
| HG02236 | ERR3242206 | IBS | Iberian populations in Spain | EUR |
| HG02224 | ERR3242251 | IBS | Iberian populations in Spain | EUR |
| HG01500 | ERR3241916 | IBS | Iberian populations in Spain | EUR |
| HG01503 | ERR3241918 | IBS | Iberian populations in Spain | EUR |
| HG00269 | ERR3240226 | FIN | Finnish in Finland | EUR |
| HG00309 | ERR3240232 | FIN | Finnish in Finland | EUR |
| HG00313 | ERR3240234 | FIN | Finnish in Finland | EUR |
| HG00121 | ERR3240195 | GBR | British in England and Scotland | EUR |
| HG00159 | ERR3240200 | GBR | British in England and Scotland | EUR |
| HG00232 | ERR3240202 | GBR | British in England and Scotland | EUR |
| HG00449 | ERR3240188 | CHS | Han Chinese South, China | EAS |
| HG00421 | ERR3241670 | CHS | Han Chinese South, China | EAS |
| HG00472 | ERR3241677 | CHS | Han Chinese South, China | EAS |
| HG02121 | ERR3242211 | KHV | Kinh in Ho Chi Minh City, Vietnam | EAS |
| HG02035 | ERR3242341 | KHV | Kinh in Ho Chi Minh City, Vietnam | EAS |
| HG02040 | ERR3242342 | KHV | Kinh in Ho Chi Minh City, Vietnam | EAS |
| HG02396 | ERR3242216 | CDX | Chinese Dai in Xishuangbanna, China | EAS |
| HG02250 | ERR3242260 | CDX | Chinese Dai in Xishuangbanna, China | EAS |
| HG02360 | ERR3242265 | CDX | Chinese Dai in Xishuangbanna, China | EAS |
| HG02380 | ERR3242272 | CDX | Chinese Dai in Xishuangbanna, China | EAS |
| HG01882 | ERR3242189 | ACB | African Caribbeans in Barbados | AFR |
| HG01886 | ERR3242194 | ACB | African Caribbeans in Barbados | AFR |

| Sample ID | ENA run accession | Population code | Population/region | Superpopulation |
|---|---|---|---|---|
| HG02012 | ERR3242223 | ACB | African Caribbeans in Barbados | AFR |
| NA18867 | ERR3239547 | YRI | Yoruba in Ibadan, Nigeria | AFR |
| NA18924 | ERR3239554 | YRI | Yoruba in Ibadan, Nigeria | AFR |
| NA19096 | ERR3239619 | YRI | Yoruba in Ibadan, Nigeria | AFR |
| HG03267 | ERR3242588 | ESN | Esan in Nigeria | AFR |
| HG03291 | ERR3242593 | ESN | Esan in Nigeria | AFR |
| HG03295 | ERR3242594 | ESN | Esan in Nigeria | AFR |
| HG03366 | ERR3242599 | ESN | Esan in Nigeria | AFR |
| HG03780 | ERR3242911 | ITU | Indian Telugu in the United Kingdom | SAS |
| HG03779 | ERR3242915 | ITU | Indian Telugu in the United Kingdom | SAS |
| HG03861 | ERR3242922 | ITU | Indian Telugu in the United Kingdom | SAS |
| HG03963 | ERR3242927 | ITU | Indian Telugu in the United Kingdom | SAS |
| HG03753 | ERR3242892 | STU | Sri Lankan Tamil in the United Kingdom | SAS |
| HG03848 | ERR3242897 | STU | Sri Lankan Tamil in the United Kingdom | SAS |
| HG03856 | ERR3242899 | STU | Sri Lankan Tamil in the United Kingdom | SAS |
| HG03490 | ERR3242630 | PJL | Punjabi in Lahore, Pakistan | SAS |
| HG03619 | ERR3242632 | PJL | Punjabi in Lahore, Pakistan | SAS |
| HG03016 | ERR3242637 | PJL | Punjabi in Lahore, Pakistan | SAS |
| HG02301 | ERR3242201 | PEL | Peruvian in Lima, Peru | AMR |
| HG02259 | ERR3242239 | PEL | Peruvian in Lima, Peru | AMR |
| HG02292 | ERR3242246 | PEL | Peruvian in Lima, Peru | AMR |
| HG01173 | ERR3241844 | PUR | Puerto Rican in Puerto Rico | AMR |
| HG01176 | ERR3241846 | PUR | Puerto Rican in Puerto Rico | AMR |

| Sample ID | ENA run accession | Population code | Population/region | Superpopulatio n |
|---|---|---|---|---|
| HG01188 | ERR3241851 | PUR | Puerto Rican in Puerto Rico | AMR |
| HG01251 | ERR3241863 | CLM | Colombian in Medellin, Colombia | AMR |
| HG01257 | ERR3241867 | CLM | Colombian in Medellin, Colombia | AMR |
| HG01357 | ERR3241879 | CLM | Colombian in Medellin, Colombia | AMR |
| HG01384 | ERR3241883 | CLM | Colombian in Medellin, Colombia | AMR |

Note: Population and superpopulation assignments follow the official 1000 Genomes Project Phase 3 sample panel. Population descriptions follow International Genome Sample Resource nomenclature.

**S7.5 WBC WES cohort**

The clinical-data benchmark used 50 de-identified whole-exome sequencing datasets generated from whole-blood-cell samples that served as matched normal controls for patients with solid tumors. The datasets were provided by Geneseeq (Nanjing, China). Tumor sequencing reads and clinical phenotype data were not included.

All samples contained 150-bp paired-end reads. Analysis was restricted to chromosomes 19 and 22.

To provide a common linear-alignment basis for the comparator tools, paired FASTQ files were aligned to the same GRCh38 chromosome 19 and chromosome 22 reference using BWA-MEM2 executable version 2.2.1. Alignments were sorted and indexed using SAMtools version 1.23. HipSTR, GangSTR, and ExpansionHunter analyzed the same BAM files, whereas STR-PG analyzed the corresponding paired FASTQ files directly.

No independent biological truth set was available for the WBC cohort. This experiment therefore evaluated cross-tool diploid genotype concordance, jointly callable genotype fractions, allele repeat-count differences, robustness to quality filtering, and STR-PG agreement with an independent majority consensus of the three comparator tools. The reported concordance metrics were not interpreted as sensitivity, specificity, or accuracy against biological truth.

In the primary strict-matching analysis, loci were considered equivalent only when chromosome, start coordinate, end coordinate, and canonicalized repeat motif were identical. A secondary sensitivity analysis allowed unique one-to-one matches with the same canonical motif and strongly overlapping or nearby repeat boundaries.

No separate capture-overlap filter was applied. A sample-locus record entered the all-valid pairwise analysis when both tools produced a parseable, non-missing diploid genotype. The two alleles were sorted before comparison, and phase was ignored. The high-confidence analysis additionally applied tool-specific quality and depth thresholds, as listed in Table S5.

For the independent comparator-consensus analysis, a consensus genotype was defined when at least two of HipSTR, GangSTR, and ExpansionHunter reported the same unordered diploid genotype. STR-PG was excluded from the voting procedure and was evaluated only against the resulting comparator consensus. Confidence intervals for summary metrics were calculated using 500 bootstrap replicates.

**S7.6 STR-PG software settings and primary parameters**

STR-PG used syncmer parameters k = 15, s = 5, and t = 2, with 100-bp flanking anchors. Pointer-locus assignment required at least two informative, chain-consistent seeds. The diagonal-chain slack was 10 bp. Paired reads supporting the same pointer received a score bonus of 0.5. A read or read pair was considered ambiguous and excluded when the difference between the best and second-best pointer scores was below 0.25.

The production-default Smith-Waterman backend used a match score of 2, mismatch score of -5, gap score of -5, alignment band of 500, and softmax temperature of 12. Informative reads were required to connect a 12-bp flank anchor to at least two adjacent motif units, with a motif mismatch fraction no greater than 0.10.

The optional pHMM backend used a mismatch probability of 0.01. Flank indel-open and indel-extension probabilities were 0.003 and 0.10, respectively; repeat-region indel-open and indel-extension probabilities were 0.005 and 0.30, respectively. The insertion-background probability was 0.25 per nucleotide. The pHMM was treated as an experimental backend and was used only in the corresponding ablation configuration rather than in the production-default results.

The optional mixture model used deterministic multi-start expectation-maximization with a maximum of 200 iterations and a convergence tolerance of $10^{-7}$. The mixture model was used as a diagnostic of within-sample repeat-length heterogeneity.

Novel-allele discovery was disabled by default. When enabled, a candidate novel allele required support from at least three reads, a motif mismatch fraction no greater than 0.10, a stutter-shadow fraction no greater than 0.15, a log-score gain of at least 3, genotype quality of at least 20, and inclusion of the candidate allele in the maximum-a-posteriori genotype.

Registry learning used count-space updates. Direct population-frequency updates were restricted to high-confidence homozygous calls. When historical allele counts were unavailable, frequency tables were initialized using a pseudo-count base total of 1000.

#### S7.7 Cross-tool software settings and genotype normalization

The WBC cross-tool concordance analysis used HipSTR version 0.7, GangSTR version 2.5.0, ExpansionHunter version 5.0.0, BWA-MEM2 executable version 2.2.1, SAMtools version 1.23, and Python version 3.11.15. GraphTyper was used in the simulated-cohort comparison, whereas vg version 1.60.0 and minigraph version 0.20 were used in the regional graph-construction and scalability comparisons.

STR-PG genotypes were obtained from the allele1 and allele2 output fields. HipSTR repeat counts were reconstructed from the reference repeat count and the GB/PERIOD representation. GangSTR and ExpansionHunter repeat counts were obtained from FORMAT/REPCN. ExpansionHunter intervals were normalized from POS + 1 through INFO/END. Allele pairs from all tools were sorted before comparison.

**Table S5 Cross-tool call definitions, quality thresholds, and genotype-normalization rules**

| Tool or analysis | All-valid definition | High-confidence definition | Genotype representation or rule |
|---|---|---|---|
| STR-PG | Parseable, non-missing diploid genotype | GQ ≥ 20 and informative depth ≥ 5 | allele1/allele2 |
| HipSTR | Sample FILTER = PASS and parseable genotype | FILTER = PASS, Q ≥ 0.90, and DP ≥ 5 | Reference repeat count plus GB/PERIOD |
| GangSTR | Non-missing REPCN | Q ≥ 0.90 and DP ≥ 5 | FORMAT/REPCN |
| ExpansionHunter | Non-missing REPCN | Site FILTER = PASS or ., and LC ≥ 5 | FORMAT/REPCN |
| Strict locus matching | Same chromosome, canonical motif, start, and end | Same rule | 1-based inclusive intervals |
| Extended locus matching | Strict matches plus unique one-to-one nearby or | Same rule | Sensitivity analysis |

| Tool or analysis | All-valid definition | High-confidence definition | Genotype representation or rule |
|---|---|---|---|
| | overlapping loci with the same canonical motif | | |
| Comparator consensus | At least two comparators report the same unordered diploid genotype | Consensus formed from calls passing tool-specific high-confidence thresholds | STR-PG excluded from voting |
| Statistical uncertainty | 500 bootstrap replicates | 500 bootstrap replicates | Confidence intervals for summary metrics |

Note: The strict exact-coordinate analysis was used for Figure 6, whereas extended locus matching was evaluated as a sensitivity analysis.

The SHA-256 checksums of the chromosome-specific comparator resources were as follows:

ExpansionHunter catalog, chr19: `35715d65f13ee0e6939aab5690b0a2924f87fe0933076e726597b6ff78184d5c`

ExpansionHunter catalog, chr22: `0e2cc327eb63602539b01a0b9ac1ac0f33ca60851799079d17db1eca77bff0ad`

GangSTR catalog, chr19: `109678a0584647ca4113df2f1fcbd929d71879594a4a5ff5db42fca9d9fc64ea`

GangSTR catalog, chr22: `4da429cd2df249bdb5b31d21e50c7f143ad0d071a7ab3e4f7bea9c277cb0dffe`

HipSTR catalog, chr19: `d325a0790cce67bf0b57f5b9f16f3944343692109b6912332947b19dd9096ec7`

HipSTR catalog, chr22: `7f49bf3bf7888e3b3762ef8780e81d3229becf8167a05efacdb4fd89bad44585`

#### S7.8 Ablation configurations

The ablation experiment included 1200 independent sample-locus records. Every configuration was evaluated using the same read evidence, candidate allele set, genotype enumeration procedure, and truth labels. The primary endpoint was unordered exact diploid genotype concordance. Novel-allele discovery was disabled in every configuration to keep the candidate-allele space constant across comparisons.

**Table S6 STR-PG ablation configurations**

| Configuration | Backend | Population frequency | BN correction | Length smoothing | Mixture diagnostic |
|---|---|---|---|---|---|
| S0_FULL_SW | SW | On | On | On | On |
| S1_UNIFORM | SW | Off | Off | Off | On |
| S2_HWE_ONLY | SW | HWE only | Off | Off | On |
| S3_NO_BN | SW | On | Off | On | On |
| S4_NO_SMOOTH | SW | On | On | Off | On |
| S5_PHMM | pHMM | On | On | On | On |
| S6_SIMPLE_SW | SW | HWE only | Off | Off | Off |
| S7_DIPLOID_ONLY | SW | On | On | On | Off |

Note: S0_FULL_SW was the production-default configuration. S1-S7 removed or replaced individual components to quantify the contributions of population-frequency information, Balding-Nichols correction, length smoothing, the mixture diagnostic, and the likelihood backend.

**S7.9 Computational environments and resource measurements**

The 1-Mb regional scalability benchmark was executed on an Intel Xeon Gold 6248R processor operating at 3.00 GHz with 512 GB RAM under CentOS 7. Runtime and peak resident memory were measured using /usr/bin/time -v.

The WBC cross-tool analysis used a Rocky Linux 8.8 software environment with Python 3.11.15. The final statistical-summary job was submitted to the p1 Slurm partition with four CPU cores, 32 GB requested memory, and a 12-h wall-clock limit. Confidence intervals were calculated using 500 bootstrap replicates.

The main operational measurements from the whole-genome experiments are summarized in Table S7.

**Table S7 Operational whole-genome resource measurements**

| Task | Dataset scale | Recorded metric | Measurement |
|---|---|---|---|
| Pointer-augmented graph construction | GRCh38 plus the stated 1000 Genomes resources | Construction time | 21.36 h |
| Pointer-augmented graph construction | GRCh38 plus the stated 1000 Genomes resources | Index size | 6.3 GB |
| Registry-level addition of one sample | One additional sample | Update time | 4.3 s |
| Pointer-level read localization | One 30× WGS sample | GAF output size | 114.23 GB |

Note: All runtime and resource measurements correspond to the stated dataset scale, software configuration, and computational setting. Runtime and memory use may vary with hardware, thread allocation, storage performance, and input-data composition.

[S9] Sehgal A, Ziaei Jam H, Shen A, Gymrek M. Genome-wide detection of somatic mosaicism at short tandem repeats. Bioinformatics 2024;40:btae485.

[S10] Ziaei Jam H, Li Y, DeVito R, Mousavi N, Ma N, Lujumba I, et al. A deep population reference panel of tandem repeat variation. Nat Commun 2023;14:6711.

[S11] Lundström OS, Verbiest MA, Xia F, Ziaei Jam H, Zlobec I, Anisimova M, et al. WebSTR: a population-wide database of short tandem repeat variation in humans. J Mol Biol 2023;435:168260.

[S12] Edgar RC. Syncmers are more sensitive than minimizers for selecting conserved k-mers in biological sequences. PeerJ 2021;9:e10805.

[S13] Eddy SR. Profile hidden Markov models. Bioinformatics 1998;14:755–63.

[S14] Wang S, Wang J, Xiao X, Zhang X, Wang X, Zhu X, et al. GSDcreator: an efficient and comprehensive simulator for genarating NGS data with population genetic information. In: Proceedings of the 2019 IEEE International Conference on Bioinformatics and Biomedicine. Piscataway: IEEE; 2019, p. 1868–75.

[S15] 1000 Genomes Project Consortium. A global reference for human genetic variation. Nature 2015;526:68–74.

[S16] Guo S, Huang Z, Zhang Y, He Y, Chen X, Wang W, et al. Enhancing variant calling in whole-exome sequencing data using population-matched reference genomes. Genomics Proteomics Bioinformatics 2024;22:qzae070.
